# Histone tails cooperate to control the breathing of genomic nucleosomes

**DOI:** 10.1101/2020.09.04.282921

**Authors:** Jan Huertas, Hans R Schöler, Vlad Cojocaru

## Abstract

Genomic DNA is packaged in chromatin, a dynamic fiber variable in size and compaction. In chromatin, repeating nucleosome units wrap 146 DNA basepairs around histone proteins. Genetic and epigenetic regulation of genes relies on structural transitions in chromatin which are driven by intra- and internucleosome dynamics and modulated by chemical modifications of the unstructured terminal tails of histones. Here we demonstrate how the interplay between histone H3 and H2A tails control ample nucleosome breathing motions. We monitored large openings of two genomic nucleosomes, and only moderate breathing of an engineered nucleosome in atomistic molecular simulations amounting to 18μs. Transitions between open and closed nucleosome conformations were driven by the displacement and changes in compaction of the two histone tails. These motions involved changes in the DNA interaction profiles of clusters of epigenetic regulatory aminoacids in the tails. Histone tail modulated nucleosome breathing is a key mechanism of chromatin dynamics.

## Introduction

In eukaryotic cells, the DNA is packed into chromatin, a dynamic fiber structure made of arrays of nucleosomes. In each nucleosome, 146 basepairs of DNA are wrapped around a protein octamer made of 4 histones (H3, H4, H2A, H2B), each present twice (*1, 2*). The N-terminus regions of all histones and the C-terminus of H2A (H2AC) are disordered, positively charged tails, carrying ~28% of the mass of the structured core of the histones (*1*).

Histone tails interact with the DNA in a non-specific manner, protruding from the nucleosome superhelix and embracing the DNA. Their role in controlling the gene expression has been extensively analyzed (*3–5*). H3 and H4 tails mediate the interaction between nucleosomes (*6, 7*), while the H2AC tail affects nucleosome dynamics, repositioning, and interacts with proteins modulating chromatin dynamics, such as linker histones or chromatin remodelers (*8*). The charged residues of histone tails are targets for “post-translational modifications”: additions of chemical groups that act as epigenetic markers to regulate chromatin accessibility and gene expression at a given time in a specific cellular context (*3*). For example, the tri-methylation of K9 (*9*) or K27 in H3 (*10*) mark regions of closed, inactive chromatin, contributing to gene silencing. Contrarily, lysine acetylation in H3 (*11*) and H4 (*12*) mark open, actively transcribed regions of chromatin. These tails modulate chromatin structure by controlling both inter- and intra-nucleosome interactions.

The impact of the tails on mononucleosome structural dynamics has been investigated with high resolution techniques such as single-molecule Förster Ressonance Energy Transfer. Such experiments confirmed the role of the H3 tail as a “close pin” that modulates nucleosome breathing and unwrapping (*13*). However, characterizing histone tail structure and dynamics at atomic resolution is difficult. Only one experimentally determined structure of nucleosomes has the histone tails resolved (*14*).

Nucleosome structure and dynamics at atomic resolution can be measured from molecular dynamics (MD) simulations. In these, time traces of the molecular system are generated by solving Newton’s equation of motion. Because of the long time scale of large amplitude nucleosome motions and the bulky, disordered nature of the histone tails, the computational resources required for atomistic simulations are prohibitive. For this reason, many simulations were performed with simplified, coarse grained nucleosomes (*15–22*). From these, the role of DNA sequence in nucleosome unwrapping (*17,19, 20*), protein-mediated nucleosome remodeling (*22*), or the impact of histone tails on nucleosome mobility, intra-nucleosome interactions and nucleosome unwrapping (*15,18,21,23*) were evaluated. However, the simplifications used in coarse grained models hinder the accurate characterization of essential physical atomic interactions.

For a complete characterization of how histone tails interact with DNA and control nucleosome dynamics all-atom MD simulations are indispensable. Such simulations have been mostly performed with engineered or incomplete nucleosomes and the sampling achieved was limited. For example, from biased MD simulations of nucleosomes without histone tails, the histone-DNA interactions involved in nucleosome unwrapping were determined (*24*). Nucleosome opening has been observed at high salt concentrations when the tails were partially truncated (*25*). The hydration patterns around the nucleosome core (*26*) and the local flexibility of nucleosomal DNA have been characterized (*27–29*). However, these studies provide only a partial description of nucleosome dynamics due to the absence of histone tails.

The structural flexibility of the tails was demonstrated in simulations of free histones or tail mimicking peptides (*30, 31*). In the few atomistic simulations of complete nucleosomes reported to date, the structural flexibility was limited due to the short time scales or the small number of trajectories (*32–38*). For example, limited nucleosome breathing and the collapse of histone tails on DNA was observed in a few 1*μ*s simulations (*35*). The mechanisms by which H2A histone variants alter nucleosome dynamics were proposed from an ensemble of 4 600 ns long simulations (*36*). From a 3.36*μ*s ensemble of ~100 ns short trajectories at high temperature (353 K), opening of truncated nucleosomes with either the H3 or H2AC tails removed was observed (*37*).

These studies have been performed using mainly two DNA sequences: a palindromic sequence derived from human *α*-satellite DNA present in the crystal structure with tails (*14*), and the Widom 601 sequence, an artificial sequence selected for strong positioning and high stability of the nucleosomes (*39*) also present in several crystal structures (*40,41*). For gene regulation in cells, the structural flexibility and mobility of nucleosomes with genomic DNA sequences matters most: genomic nucleosomes are more dynamic than engineered ones (*42*). Recently, we reported large amplitude breathing and twisting motions from MD simulations of 2 genomic nucleosomes (*38*), which bear enhancer sequences of the genes *ESRRB* and *LIN28B.* These genes are important for stem cell pluripotency (*43, 44*) and their nucleosome wrapped enhancers are bound by the key pioneer transcription factor Oct4 when converting skin to stem cells (*45,46*). Our simulations were aimed at revealing the binding modes of Oct4 to these nucleosomes.

Here, from a total of 18 *μ*s atomistic simulations of the engineered Widom and the genomic Lin28b and Esrrb nucleosomes, including the already reported 9 *μ*s performed with *Drosophila* histones (from now on dH simulations) and additional 9 *μ*s with human histones (hH simulations), we demonstrate how the interplay between histone H3 and H2AC tails controls nucleosome breathing motions. We monitored 2 rare major opening events and found that nucleosome conformations with different degree of opening display specific H3 and H2AC conformations and positions. Moreover, nucleosome opening and closing is regulated by specific patterns of interactions between epigenetic regulatory residues in H3 and H2AC tails and the DNA. Because the distribution of open and closed nucleosomes impacts their structure and compaction of chromatin fibers, the mechanism we describe is key to understanding chromatin dynamics.

## Materials and Methods

### Nucleosome Modeling

The 146 bp Widom nucleosomes with *Drosophila melanogaster* and Homo Sapiens histones were built by homology modelling, using Modeller (https://salilab.org/modeller/), with the same procedure described in our previous report (*38*). As templates, we used the structure of the *Drosophila melanogaster* nucleosome core (PDB ID: 2PYO), the structure of the Widom 601 nucleosome particle (PDB: 3LZ0), and the crystal structure of the X. laevis nucleosome that includes histone tails (PDB: 1KX5). For each nucleosome, 100 homology models were generated using a “slow” optimization protocol and a “slow” MD-based refinement protocol as defined in Modeller. We selected the lowest energy model and added an 11 bp fragment of B-DNA (generated with Nucleic Acid Builder from Ambertools 18 (*47*)) to each linker DNA to generate the 168 bp nucleosomes.

The Esrrb and Lin28b enhancer sequences were the same as used in our previous work (*38*) (Supplementary Document S1), originally selected from data by Soufi et al (*45*). We substituted the Widom sequence with the Lin28b and the Esrrb sequences using the “swapna” function in Chimera (https://www.cgl.ucsf.edu/chimera/). In the human Lin28b nucleosome, the sequence was shifted by 1 extra bp (2 compared to the construct presented in Soufi et al. (*46*)). This shift was introduced following our previous work (38) to optimize the accessibility of the Oct4 binding sites.

### Molecular dynamics simulations

The MD protocol was described previously (*38*). In short, each nucleosome was solvated in a truncated octahedron box of at least 12 Å of SPCE water molecules around the solute in any direction. Na^+^ ions were added to neutralize the charge of the system, and K^+^ and Cl^-^ ions were added up to a concentration of 150mM KCl. Then an energy minimization using the AMBER software (*47*) was performed, followed by 10.25 ns of equilibration with gradually decreasing positonal and base pairing restraints (*38, 48*). The equilibration and the following production runs were performed using NAMD (*49*). The production runs were in the isobaric-isothermic (NPT, p = 1 atm T = 300 K) ensemble, using Langevin dynamics as a termostat and Nosé-Hoover and Langevin piston for pressure control. The AMBER force field parameters ff14SB (*50*), parmbsc1 (*51*) and Li-Merz (*52*) were used for protein, DNA and ions, respectively.

### Analysis of nucleosome dynamics

All simulations were fitted with a root mean-square fit of the heavy atoms of the histone core to the minimized structure of the Widom nucleosome. The histone core was defined by excluding the histone tails: residues 1—45 for hH3 and dH3, 1—32 for hH4 and dH4, 1-18 and 129 for hH2A, 1—17 and 115—124 for dH2A, and 1-33 for hH2B and 1—31 for dH2B.

The characterization of the breathing motions was done using the procedure first described by Ozturk et al. (*53*), that we also used in our previous work (*38*). We first defined a coordinate system *XYZ* with the origin on the dyad. *X* was defined as the vector along the dyad axis.*Y* was defined as the cross product between *X* and a vector perpendicular to *X* intersecting it approximately at the center of the nucleosome. Finally *Z* was defined as the cross product between *X* and *Y*. Then the vectors *v_3_* and v_5_ were defined along the 3’ and 5’ L-DNAs. The angle *γ*_1_ was defined as the angle between the projector of these vectors in the *XZ* plane and the *Z* axis, whereas *γ*_2_ was defined as the angle between the projection of the vectors on the *XY* plane and the *Y* axis.

The radius of gyration of the DNA (RoG) and the number of protein-DNA contacts were calculated using the cpptraj software (*54*). Contacts were defined as protein non-hydrogen atoms closer than 4.5 Å. We split the DNA in two parts. The outer gyre was defined as 40 bp from each end of the nucleosomal DNA, and the inner gyre was defined to include all remaining base pairs. For the histones, contacts were split between contacts made by the histone core and the histone tails, using the core and tail definition described above.

Mean and minimal distances were measured using the weighted mean distance collective variable *distanceInv* implemented in the Colvar module (*55*) available in VMD (*56*). The weighted distance between two groups of atoms is defined as follows: 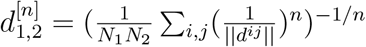, where ||*d^ij^*|| is the distance between atoms *i* and *j* in groups 1 and 2 respectively, and *n* is an even integer. This distance will asymptotically approach the minimal distance when increasing *n*. For smaller values of *n*, weighted distance will be close to the true average distance. For measuring the position of the histone tails relative to DNA, we calculated the weighted mean distances with *n* =10 (from now on referred to as “mean distances”) from the tails to two reference points: the L-DNA, defined as the last 15 basepairs of each DNA arm, and the dyad, defined as the 8 bp next to the dyad on the same side as the tail (3’ and 5’). Minimal distances between individual residues in histone tails and the inner gyre or the outer gyre of DNA (as defined above), were calculated as weighted mean distances with *n* = 100.

Principal Component Analysis (PCA) was performed using cpptraj. Due to the high flexibility of the histone tails, all histone tails except for the H3 and H2AC tails were removed for the PCA calculation. Non-hydrogen atoms of the DNA and histones (excluding the H4, H2B and H2AN tails) were used. For both the Esrrb^hH^ and the Lin28^dH^ nucleosomes, the covariance matrix was calculated and diagonalized to extract the first 25 eigenvectors and eigenvalues. The trajectories were projected on the first mode, and the minimum and maximum projection values for that mode were extracted. Finally, pseudotrajectories along the first mode were generated to analyze the correlated motions of the L-DNA and tails. PC1 represents 42.93% and 31.3% of all the motions in the Esrrb^hH^ and Lin28^dH^ respectively.

### Clustering of MD trajectories

MD trajectories of all nucleosomes with the same histones were combined, obtaining 2 groups of simulations: one with human histones (hH simulations) and one with *Drosophila* histones (dH simulations). hH and dH simulations were separated because a clustering of the combined ensemble of simulations based on the histone tails would not be possible due to the differences in histone sequences between the 2 organisms. The snapshots from each group of simulations were clustered based on the positional RMSD of the backbone heavy atoms of DNA, H3 and H2A. For the DNA, the outer gyre that didn’t display big opening events (5’ and 3’ in hH and dH simulations respectively) was not considered in the clustering. For the histone H3, the RMSD of the entire histone was used for clustering, whereas for the H2A, the RMSD of the entire histone excluding the N-terminal tail was used. This resulted in 3 independent clustering distributions, referred to as “DNA clusters”, “H3 clusters”, and “H2AC clusters”. The k-means clustering algorithm implemented in cpptraj was used, with 8 centroids. Finally, the percentage of snapshots from each DNA cluster in each H3 and H2AC cluster was computed, obtaining the distributions of the DNA cluster snapshots in the histone tail clusters to evaluate the correlation between DNA conformations and the H3 and H2AC conformations and positions.

## Results

### Breathing motions are more extensive in genomic nucleosomes

We performed 3 independent 1 *μ*s long simulations for each combination of DNA and histones (Table 1), which we grouped in ensembles of 3 *μ*s for analysis. The highest structural flexibility we monitored in the L-DNA arms (both 5’ and 3’) and the histone tails (Figure 1), shown by the broader range of the number of contacts with DNA formed by the histone tails compared to the histone core (Table1, Supplementary Figure S1). All tails interacted with the core nucleosomal DNA (146 bp centered on the dyad), but only the H3 and H2AC tails interacted also with the linker DNA (L-DNA).

**Figure 1:**
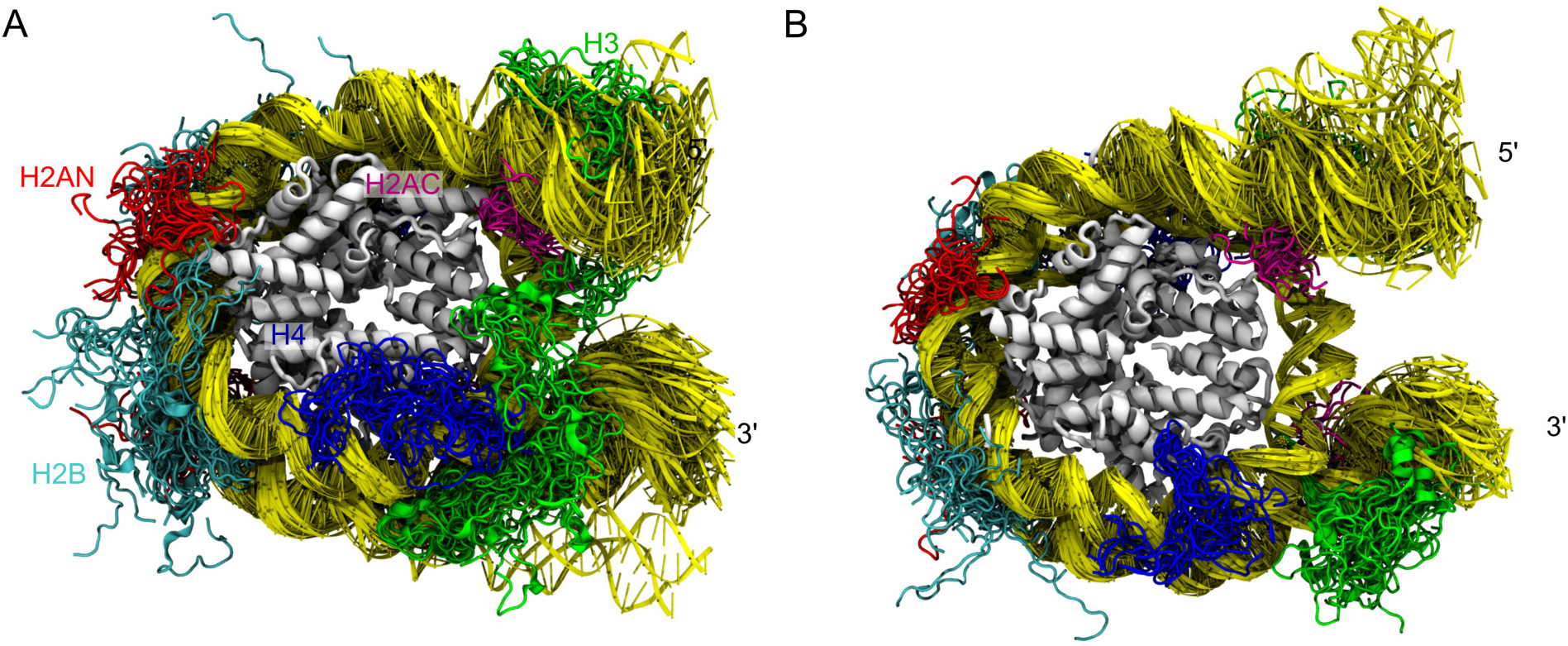
Nucleosome motions on the *μ*s time scale. Superposition of nucleosome snapshots taken from the 3 *μ*s simulation ensembles in which the major opening events occurred. **A**) Esrrb^hH^ **B**) Lin28b^dH^. Tails are shown every 100ns, the DNA every 50 ns. The histone core is in gray, DNA in yellow. The histone tails of H3, H4, H2A (both N and C terminal) and H2B are in green, blue, red, magenta and cyan respectively.

**Table 1:** Overview of the simulations performed

| Histones | DNA | Replica | Time | RoG<sup>1</sup> | Number of contacts<sup>2</sup> with DNA |  |  |
| --- | --- | --- | --- | --- | --- | --- | --- |
|  |  |  |  |  | Histone Core | Histone Tails |  |
|  |  |  |  |  |  | inner gyre<sup>3</sup> | outer gyre<sup>3</sup> |
| *Drosophila* | Widom | 1 | 1 μs | 47.11–48.34 | 1914–2124 | 790–1514 | 891–1360 |
| *Drosophila* | Widom | 2 | 1 μs | 47.62–49.58 | 1837–2151 | 683–982 | 743–1250 |
| *Drosophila* | Widom | 3 | 1 μs | 48.08–49.03 | 1780–2019 | 769–1299 | 907–1421 |
| *Drosophila* | Esrrb | 1 | 1 μs | 47.37–48.82 | 2010–2265 | 937–1388 | 764–1187 |
| *Drosophila* | Esrrb | 2 | 1 μs | 47.26–48.34 | 1869–2161 | 675–1269 | 989–1295 |
| *Drosophila* | Esrrb | 3 | 1 μs | 47.11–48.41 | 2013–2241 | 894–1375 | 783–1146 |
| *Drosophila* | Lin28b | 1 | 1 μs | 48.00–49.91 | 1917–2170 | 837–1353 | 942–1354 |
| *Drosophila* | Lin28b | 2 | 1 μs | 48.34–50.90 | 1928–2173 | 755–1718 | 639–1185 |
| *Drosophila* | Lin28b | 3 | 1 μs | 47.63–49.26 | 1977–2171 | 917–1384 | 812–1066 |
| Human | Widom | 1 | 1 μs | 47.93–49.26 | 2017–2224 | 762–1260 | 754–1031 |
| Human | Widom | 2 | 1 μs | 47.26–48.54 | 1867–2116 | 1061–1673 | 717–1023 |
| Human | Widom | 3 | 1 μs | 48.06–49.18 | 1807–2047 | 856–1442 | 716–1226 |
| Human | Esrrb | 1 | 1 μs | 47.83–51.32 | 1708–2057 | 1147–1893 | 457–763 |
| Human | Esrrb | 2 | 1 μs | 48.22–49.87 | 1844–2062 | 888–1631 | 777–1184 |
| Human | Esrrb | 3 | 1 μs | 47.14–48.18 | 1926–2134 | 901–1438 | 666–1098 |
| Human | Lin28b | 1 | 1 μs | 47.79–48.93 | 1907–2121 | 752–1123 | 828–1488 |
| Human | Lin28b | 2 | 1 μs | 47.93–49.00 | 1852–2048 | 827–1284 | 830–1261 |
| Human | Lin28b | 3 | 1 μs | 47.90–49.62 | 1792–2048 | 667–1549 | 739–1119 |

<sup>1</sup> RoG and the number of contacts are shown as the percentiles 5 to 95.
<sup>2</sup> A contact was defined as a non-hydrogen atom closer than 4.5 Å to another non-hydrogen atom.
<sup>3</sup> Inner and outer gyre of the nucleosomal DNA, see Methods for details.

To characterize structural flexibility, we computed the radius of gyration (RoG) and two angles, *γ*_1_ and *γ*_2_. These provide a complete description of nucleosome breathing in the XZ (*γ*_1_) and XY (*γ*_2_) planes defined by a dyad-centered coordinate system (*38,53*) (Figure 2A, see Methods). The main RoG peaks of the nucleosomes with human and *Drosophila* histones (hH and dH respectively) overlap, indicating a similar conformational space sampling in the 2 simulation sets. The genomic nucleosomes Lin28b and Esrrb adopted more open conformations with the exception of the Esrrb^dH^ nucleosome (Figure 2B, Supplementary Figure S2). In contrast, both Widom nucleo-somes sampled mainly conformations with RoG around ≈ 48.25 Å and less often conformations with RoG ≈ 47.5 Å. The latter was similar with the closed conformation sampled by Esrrb^dH^. The Esrrb^hH^ nucleosome sampled three regions of the conformational space, two of them have RoG values resembling those sampled by the Widom nucleosomes, while the third has more open conformations (RoG ¿50). The Lin28b nucleosomes were more open (RoG 48.25 – 49.5 Å) and did not sample the most closed conformations. Additionally, the Lin28b^dH^ sampled open conformations with RoG ¿ 50 Å, similar to those sampled by Esrrb^hH^ (Figure 2B).

**Figure 2:**
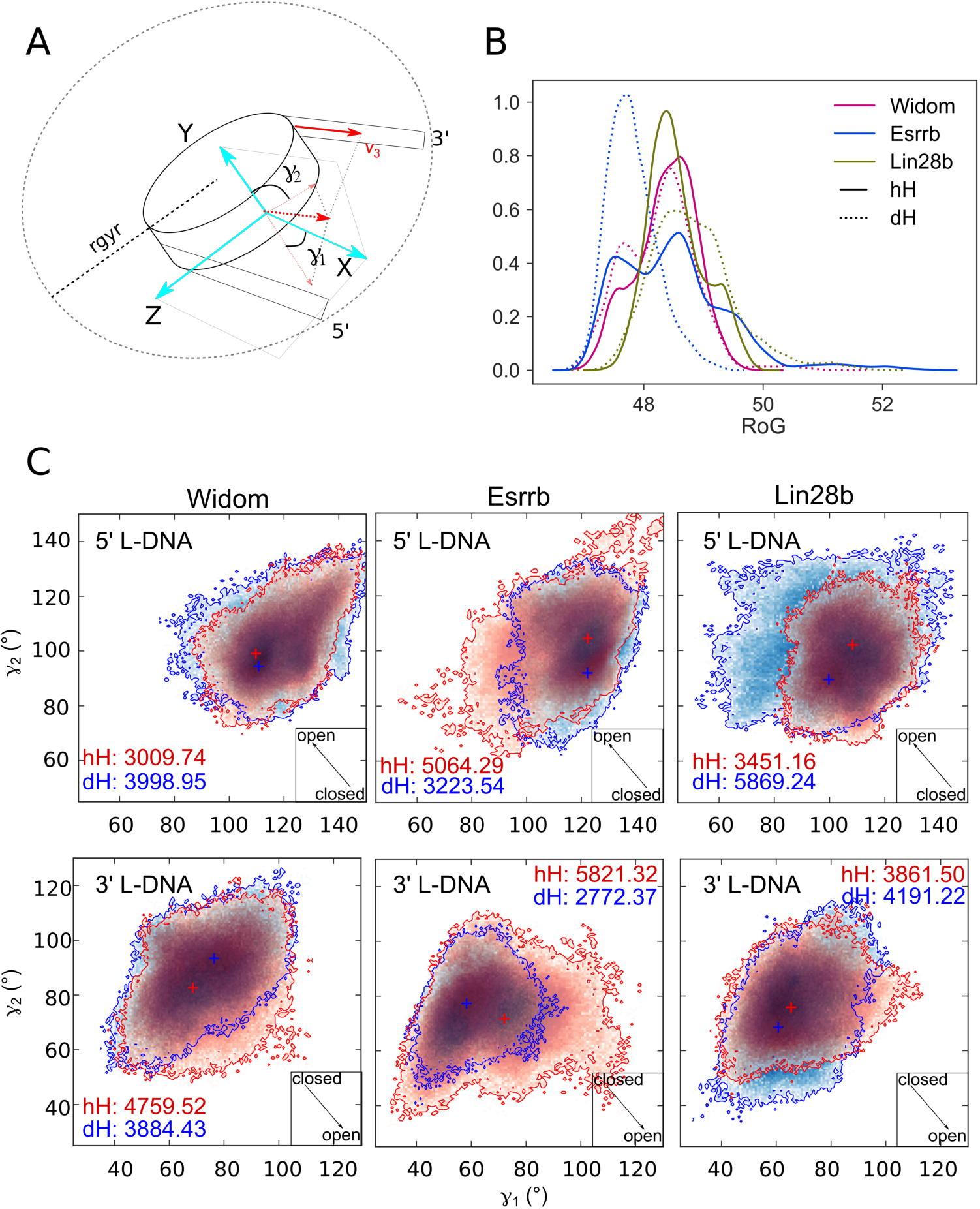
Structural flexibility of the hH and dH nucleosomes. **A)** Schematic representation of the RoG definition and the coordinate system used to describe nucleosome conformations. The angles *γ*_1_ and *γ*_2_, describing the nucleosome breathing motions were defined as described in Methods. An increase of *γ*_1_ indicates opening at the 3’ L-DNA, but closing at the 5’ L-DNA, and vice-versa for *γ*_2_. **B)** RoG distribution from the 3 *μ*s ensembles of simulations. Larger values indicate a lower degree of compaction in the nucleosomal DNA. Solid and dashed lines represent the hH and dH simulations, respectively. The Widom, Esrrb, and Lin28b nucleosomes are shown in magenta, blue and green respectively. **C)** Two-dimensional histograms depicting the conformational sampling of the 5’ and 3’ L-DNA arms in the space defined by the *γ*_1_ and *γ*_2_ angles. The arrow inserts in the lower-right corner indicate the direction of the opening, whereas the numbers are the areas of the histograms. The crosses indicate the most populated region of each histogram. The data for the hH and dH nucleosomes are in red and blue respectively.

In summary, the genomic nucleosomes breath more extensively than engineered nucleosomes on *μ*s time scale. We monitored 2 large amplitude transient openings, one at the 3’ L-DNA of Esrrb^hH^ and one the 5’ L-DNA of Lin28b^dH^.

The open conformations of Esrrb^hH^ and Lin28b^dH^ with highest RoG were sampled in simulations 2 and 1 of each ensemble respectively and were characterized by fewer contacts between the histone tails and the outer DNA gyre (Table 1, Supplementary Figure S1). The opening was either in both XZ and XY planes, or only in one plane and was induced by motions of the 3’ and 5’ L-DNA in the 2 nucleosomes respectively (Figure 2C).

### Histone H3 and H2A tails sample a broad range of configurations

To test whether the conformational plasticity and positional fluctuations of the histone tails modulate nucleosome breathing, we first analyzed the histone tail configurations, defined as the combination between the tail position and its conformation. We monitored the position of the H3 and H2AC tails relative to to the neighbouring L-DNA and the DNA segment around the dyad and the number of contacts of these tails with the inner and outer DNA gyre (see Methods).

We found that the H3 and H2AC tails neighboring the highly flexible L-DNA arms (3’ L-DNA in Esrrb^hH^ and the 5’ L-DNA in Lin28b^dH^) sampled a broad range of configurations (Figure 3, Supplementary Figure S3, Supplementary Document S3). Different configurations were sampled in different simulations and transitions between configurations occurred in the individual 1 *μ*s simulations. Moreover, specific tail configurations associate with particular conformations of the nucleosomal DNA. When the H3 tail was in an intermediate position between the L-DNA and the dyad, it acted as a bridge keeping the nucleosome closed. When the tail had a large number of contacts to the outer gyre, and a short distance to the L-DNA, the nucleosomal DNA was partially closed. When the H3 or H2AC tails were near the dyad and away from the L-DNA forming only few contacts to the outer gyre, the nucleosomes opened.

**Figure 3:**
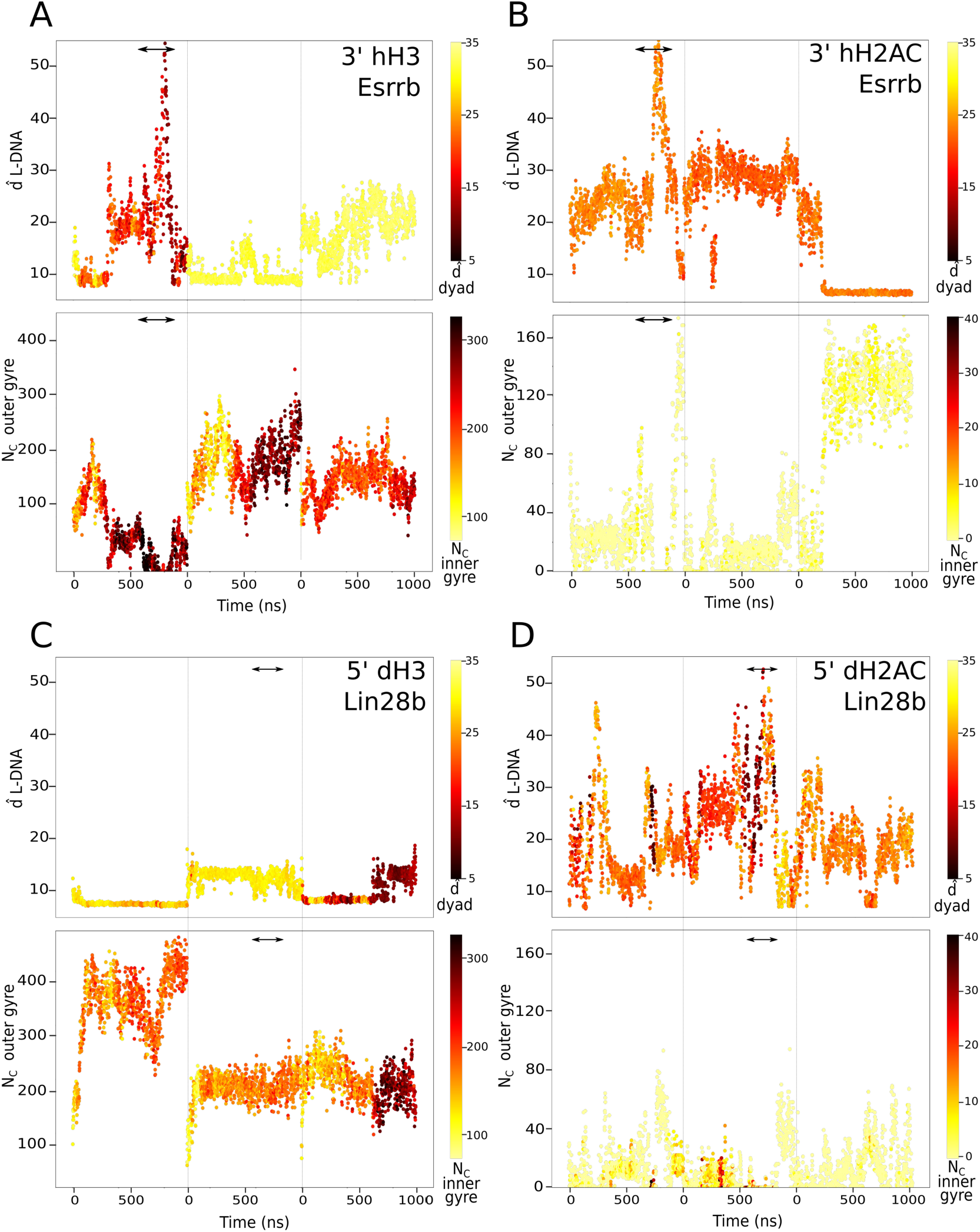
Configurations of the H3 and H2AC tails. Each panel shows the weighted mean distance to the L-DNA (top) and the number of contacts with the outer gyre (bottom) during the simulations. The points are colored by the weighted mean distance to the dyad (top) and the number of contacts to the inner gyre of the DNA (see Methods for details on the definition of inner and outer gyre). **A-B)** H3 (A) and H2AC (B) tails at the 3’ end in the simulations of the Esrrb^hH^ nucleosome. **C-D)** H3 (C) and H2AC (D) tails at the 5’ end in the simulations of the Lin28b^dH^ nucleosome

The Esrrb^hH^ nucleosome opened (750-850 ns of simulations 1) when the distance between the H3 tail and the L-DNA as well as the number of contacts of the tail with the dyad region increased. At the same time, the H2AC tail had no contacts with L-DNA while its position relative to the dyad remained stable. (Figure 3B). Towards the end of the simulation, these contacts were quickly reformed, closing the DNA. Therefore, the collapse of the H3 tail on the DNA around the dyad together with the loss of the contacts of both tails with L-DNA permitted a large nucleosome opening (Figure 3A).

The Lin28b^dH^ nucleosome opened (700-800 ns, simulation 2), when the H3 tail collapsed around the middle of the 5’ L-DNA, far from the dyad (Figure 3C). Before the opening, the configuration of the H2AC changed: its distance to the L-DNA increased while the number of contacts to the outer gyre decreased(Figure 3D). Again, the contacts with DNA reformed shortly after opening, closing the DNA.

In conclusion, the H3 and H2AC tails sampled a broad range of configurations, but only an interplay between specific positions and conformations of both tails facilitated nucleosome breathing.

### Nucleosome conformations display specific histone tail configurations

Next, we tested whether there is a systematic correlation between the H3 and H2AC tail configurations and the nucleosome conformations. We split the hH and dH simulations in 2 separate groups and clustered the snapshots from each group in 8 clusters ranked by size in 3 different ways. First, to separate open and closed nucleosome conformations, we clustered based on the RMSD of the DNA backbone in the inner gyre and the outer gyre that opens (DNA clusters) (see Methods). Second, to separate histone tail configurations, we clustered based on the RMSD of the non-hydrogen atoms of the H3 and H2A histones separately (excluding the N terminal tail of H2A) (H3 and H2AC clusters).

Then, we mapped the DNA clusters on the 2D histograms of the 7 angles (Figure 4A,B, Supplementary Figure S4) and confirmed that they spread across the sampled conformational space of the nucleosomes. The distributions of the DNA clusters from the simulations of hH and dH nucleosomes were symmetric (Figure 4A,B, Supplementary Figure S3), demonstrating that they provide a similar phase space sampling. The largest DNA clusters (*DNA*^1*h*^, *DNA*^1*d*^) contained the more closed conformations in both simulation groups, whereas the smaller clusters (*DNA*^7*h*^, *DNA*^8*h*^ and *DNA*^6*d*^) contained conformations open in both planes defined by *γ*_1_ and *γ*_2_.

**Figure 4:**
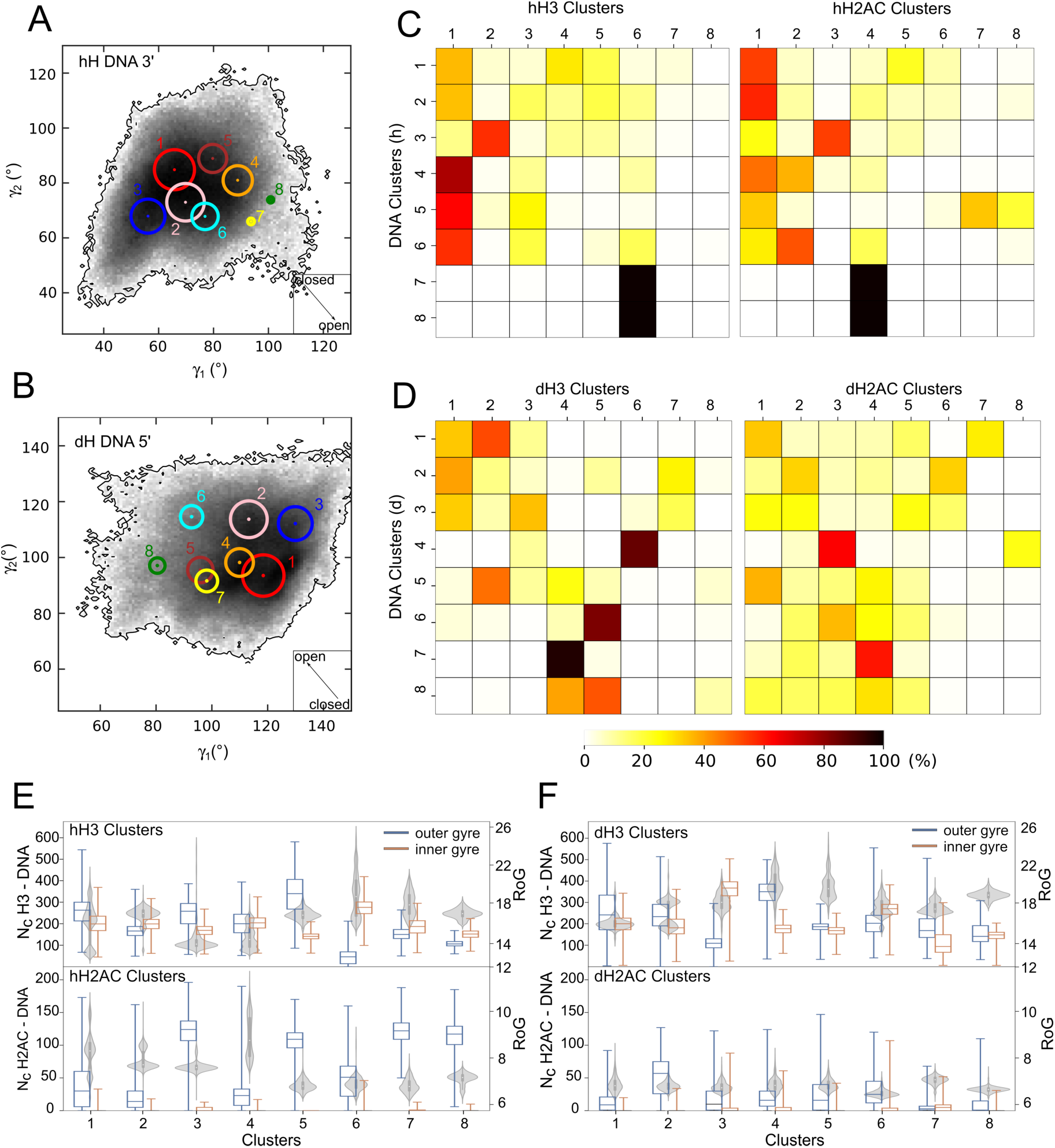
Clustering of the DNA and histone tails. **A-B**) Position of the DNA cluster centroids in the 2D histogram of the 7 distribution for the 3’ L-DNA in hH simulations (A) and 5’ L-DNA in dH simulations (B). The size of the circles indicates the size of the clusters. **C-D)** Distribution of the frames from each DNA cluster on the tail clusters (H3 in the left matrix, H2AC in the right), for hH (C) and dH (D) simulations. For each DNA cluster (row), the cells are colored by the percentage of frames of that cluster that belong to the tail cluster (column). **E-F)** Features of H3 and H2AC configurations per cluster in the hH (E) and dH simulations (F). Box plots indicate the number of contacts between H3 (top) or H2AC (bottom) tail and the inner (red) and outer gyre (blue) of DNA per tail cluster. Violin plots represent RoG of the H3 (top) or H2AC (bottom) tail per cluster.

To evaluate how DNA conformations associate with H3 and H2AC configurations, we computed the percentage of frames of each DNA cluster that is found in each H3 and H2AC cluster (Figure 4 C,D). We characterized the histone tail configurations with the RoG of the tails and the number of contacts with the inner and outer gyre of the nucleosome (Figure 4 E,F).

The most open conformations of the DNA displayed specific configurations of both H3 and H2AC tails (Figure 4C,D). In the hH simulations, the open DNA conformations from cluster *DNA*^7*h*^ (Figure 5A) and *DNA*^8*h*^ had tail configurations from clusters *hH*3^6^, and *hH2AC*^4^ (Figure 4C) with few contacts with the outer DNA gyre, a wide distribution of conformations (wide RoG distribution) and the most extended tail conformations sampled (Figure 4E). In the dH simulations, the open DNA conformations in cluster *DNA*^6*d*^ (Figure 5B) had H3 configurations from cluster *dH*3^5^, and often H2AC configurations from cluster *dH2AC*^3^ (Figure 4D). Again, *dH*3^5^ contains configurations with few contacts with both inner and outer DNA gyres and extended conformations (higher RoG) (Figure 4F). The dH2AC tail configurations are similar between different clusters because dH2AC is shorter and less positively charged compared to hH2AC (Supplementary Document S2). *dH2AC*^3^ contains configurations with a slightly higher number of contacts with the inner DNA gyre. Therefore, the large opening of both genomic nucleosomes occurred when the H3 and H2AC tails formed few interactions with the L-DNA.

**Figure 5:**
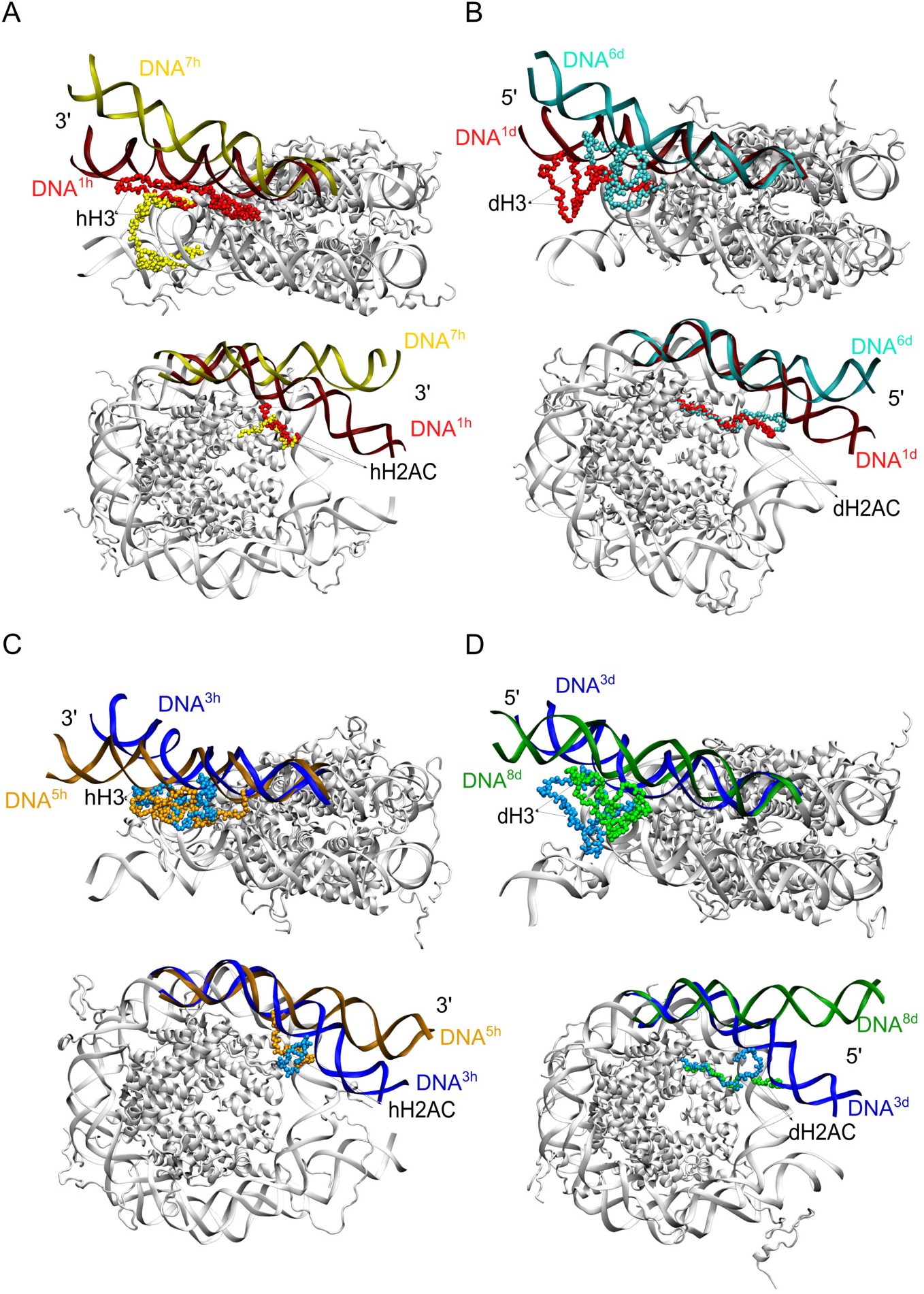
Structures of DNA cluster representatives. The L-DNA used for clustering is colored by the cluster color from Figure 4A-B. The L-DNA is in a dark shade, whereas the nearby histone tail is in a lighter shade of the same color. Each panel shows a side view of the nucleosome with the H3 tail (top), and a top view of the nucleosome with the H2AC tail (bottom). **A-B)** Conformations closed or open in both the XZ (*γ*_1_) and XY planes (*γ*_2_). For the hH simulations (A), clusters *DNA*^1*h*^ (closed) and *DNA*^7*h*^ (open). For the dH simulations (B), clusters *DNA*^1*d*^ (closed) and *DNA*^6*d*^ (open). **C-D)** Conformations open only in one plane, closed in the other. For the hH simulations (C), clusters *DNA*^3*h*^ (open in the XY plane) and *DNA*^5*h*^ (open in the XZ plane). For the dH simulations (D), clusters *DNA*^3*d*^ (open in the XY plane) and *DNA*^8*d*^ (open in the XZ plane). For clarity, the 3’ end of the hH nucleosomes is shown in the same direction as the 5’ end of the dH nucleosomes (views are inverted by 180° around the dyad axis).

The DNA conformations open only in one direction (defined by one *γ*) also displayed specific histone tail configurations. In the hH simulations, the DNA conformations open only in the direction defined by *γ*_1_ from clusters *DNA^4h^* and *DNA*^5*h*^ (Figure 5C) had almost exclusively H3 tail configurations from clusters *hH*3^1^ and *hH*3^3^ (Figure 4C) with a high number of contacts to the outer DNA gyre, and compact conformations (low RoG values) (Figure 4E). Therefore, when the nucleosome is closed in the XY plane (*γ*_2_), the H3 tail adopts a compact, bridging configuration that interacts with both DNA gyres (Figure 4E). The DNA conformations open only in the XZ plane (*γ*_1_) had H2AC configurations from the largest cluster, *hH2AC*^1^ (Figure 4C) with less contacts to the outer DNA gyre (Figure 4E). The differences between the DNA conformations in clusters *DNA*^4*h*^ and *DNA*^5*h*^ are due to their different hH2AC tail configurations found in clusters *hH2AC*^2^ and *hH*2*AC*^7^-*hH*2*AC*^8^ (Figure 4C).

The hH nucleosome conformations open only in the XY plane (*γ*_2_) (*DNA*^3*h*^, Figure 5C) had histone tail configurations from clusters *hH*3^2^ and *hH*2*AC*^3^ (Figure 4C) with extended conformations (Figure 4E). The H3 tail configurations in cluster *hH*3^2^ do not vary in their position, displaying a narrow distribution of the number of contacts with both DNA gyres (Figure 4E).

The dH nucleosomes with conformations open only in the XZ plane (*γ*_1_) in cluster *DNA*^8*d*^ (Figure 5D) had H3 configurations from clusters *dH*3^4^ and *dH*3^5^, and variable H2AC configurations. *dH*3^5^ contains tail configurations found in the most open nucleosome conformations. *dH*3^4^ contains configurations with extended conformations and more contacts with the outer DNA gyre than those in *dH*3^5^. These are present in the slightly more closed DNA conformations from cluster *DNA*^7*d*^ (Figure 4D). Therefore, the conformations in *DNA*^8*d*^ can be seen as transient between the more closed conformations in *DNA*^7*d*^ and the most open conformations in *DNA*^6*d*^. The dH nucleosome conformations open only on the XY plane (*γ*_2_) in cluster *DNA*^3*d*^ (Figure 5D) had mostly H3 configurations from clusters *dH*3^1^ and *dH*3^3^ and variable dH2AC configurations (Figure 4D). The *dH*3^3^ has more extended dH3 tail conformations with more contacts with the core nucleosomal DNA (Figure 4F). The *dH*3^1^ cluster is the largest dH3 cluster with more diverse dH3 configurations (Figure 4F). These findings further confirm that the loss of a large fraction of the contacts between H3 and H2AC tails and the L-DNAs is required for nucleosome opening.

The most closed DNA conformations in the clusters *DNA*^1*h*^, *DNA*^1*d*^ (Figure 5A) display H3 tail configurations found in the largest H3 clusters or distributed among different clusters (Figure 4C,D) that are either extended or more compact and have a large number of contacts both with L-DNAs and the core DNA (Figure 4E,F). Therefore, the nucleosomes are maintained in a closed conformation by H3 tail mediated bridging interactions between the L-DNA and the core nucleosomal DNA. The closed conformations of the hH nucleosome have H2AC tail configurations with extended conformations and a moderate number of contacts with the L-DNA (Figure 4C,E). In contrast, those of the dH nucleosomes do not display any specific H2AC tail configurations(Figure 4D,F)

The clustering analysis demonstrated that the open and closed nucleosome conformations are characterized by specific H3 and H2AC tail conformations and positions. In particular open nucleosome conformations have H3 and H2AC tail configurations with fewer contacts with the DNA, especially with the adjacent L-DNA.

### Nucleosome opening is induced by H3 and H2AC tail dynamics

To test whether the motions of the H3 and H2AC tails are a consequence of nucleosome breathing or induce it, we mapped the structural snapshots from the DNA, H3, and H2AC clusters on the simulation time traces, and compared the distributions with the nucleosome breathing profiles given by the time evolution of the RoG (Figure 6).

**Figure 6:**
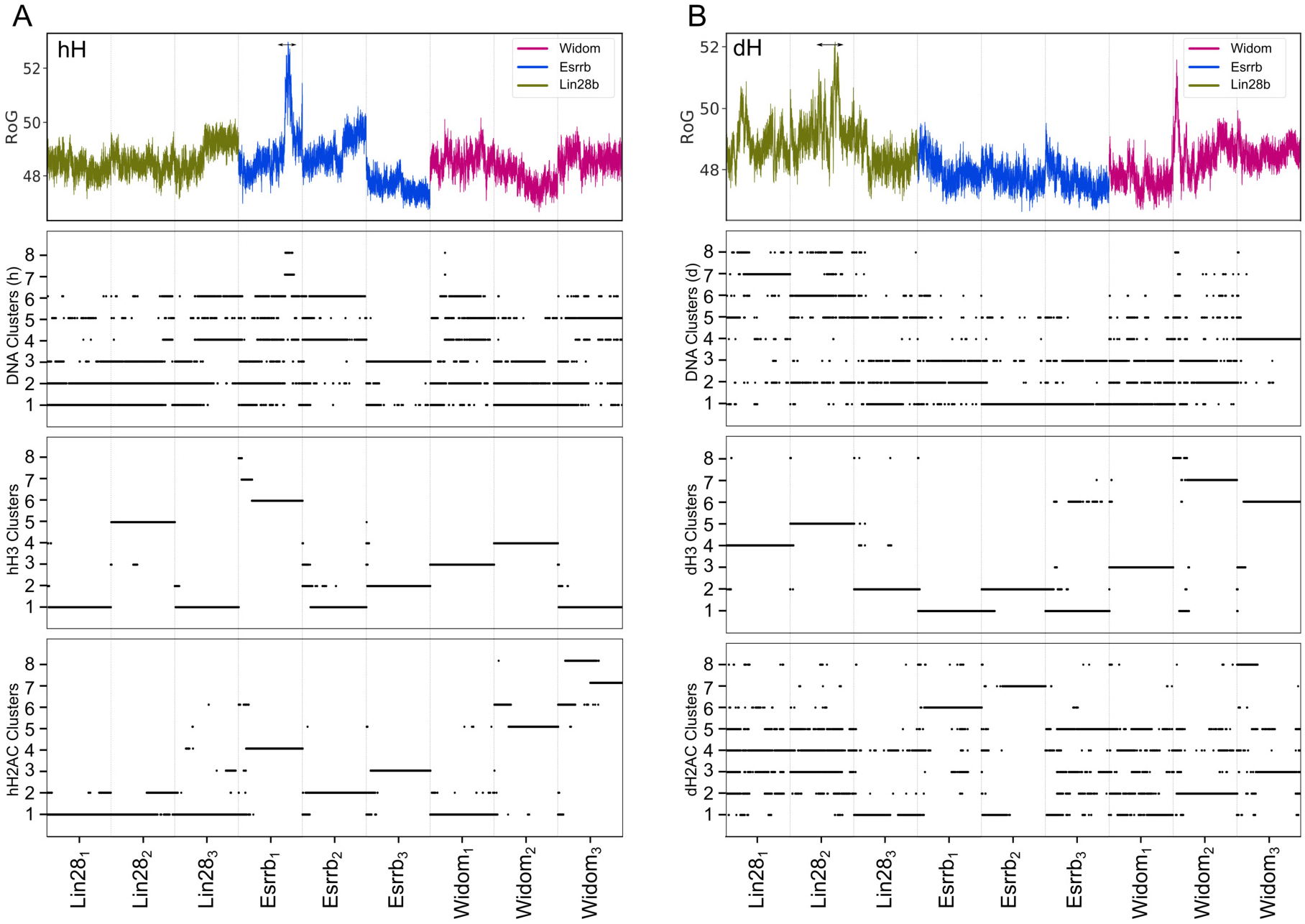
Structural transitions in the DNA, H3 and H2AC tails. Structures from each DNA, H3, and H2AC cluster mapped on the simulation time series. **A**) The 3’ end of the hH nucleosomes. **B**) The 5’ end of the dH nucleosomes. In each panel, the top plot shows the nucleosome RoG and the 3 plots below show to which DNA, H3, and H2AC clusters each simulation snapshot belongs to. Each point corresponds to one single snapshot.

We found that reversible conformational transitions in the nucleosomal DNA are frequent in 1 *μ*s, while transitions to largely open conformations are rare. In the hH simulations, the most open DNA conformations from clusters *DNA*^7*h*^ and *DNA*^8*h*^ are sampled only in the second simulation of the Esrrb^hH^ nucleosome between 700 and 800 ns (Figure 6A). In the dH simulations, they are in clusters *DNA*^6*d*^ and *DNA*^8*d*^ and are sampled mainly in the first and second simulations of the Lin28b^dH^ (Figure 6B). The opening of Lin28b^dH^ was more gradual than that of Esrrb^hH^.

In contrast, transitions between configurations of the H3 tail were rare in 1 *μ*s (Figure 6), different configurations being sampled in the independent simulations. This highlights the importance of grouping the independent simulations in ensembles for investigating such large scale motions. Transitions between H2AC tail configurations in 1 *μ*s were more frequent, especially for the shorter dH2AC.

Both large openings of the nucleosomal DNA were preceded by transitions of both H3 and H2AC tail configurations. Esrrb^hH^, opened after both H3 and H2AC tails adopted specific conformations from clusters *hH*3^6^ and *hH*2*AC*^4^ (Figure 6A). Lin28b^dH^ opened when the H3 tail was in a configuration from the *dH*3^5^ cluster and the H2AC tail transited to configurations from clusters *dH*2*AC*^6^ and *dH*2*AC*^8^ before the nucleosome opened. We confirmed the correlation between nucleosome opening and histone tail dynamics by a principal component analysis of the simulation ensembles in which the nucleosome opened (Supplementary Figure S5)

The interplay between both tails also contributed to a very short lived opening in the first simulation of the Widom^dH^ nucleosome (Table 1, Figure 6B). Here, the H3 tail sampled configurations of the *dH*3^3^ cluster with less contacts to the outer DNA gyre, and higher RoG values indicating a more extended conformation (Figure 4F). However, because the dH2AC tail didn’t adopt configurations permissive for DNA opening (*dH2AC*^6^ and *dH*2*AC*^8^) the nucleosome remained mostly is a very closed conformations.

In conclusion, the extensive opening of the nucleosomes developed only after both the H3 and H2AC tails adopted specific configurations lacking the interactions required to maintain the nucleosomes closed. Transition out of these configurations lead to nucleosome closing.

### Nucleosome opening is modulated by epigenetic regulatory residues

To test how how interactions between aminoacids in the histone tails and the DNA impact on nucleosome conformational dynamics, we monitored the minimal distance between positively charged lysines and arginines and the inner and outer DNA gyres. (Figure 7). Post-translational modifications of these key residues mark active versus inactive chromatin, are involved in epigenetic regulation of gene expression, and are expected to impact on nucleosome dynamics. We separated the residues of the H3 tail in 4 groups: the “tip” group includes the residues at the tip of the tail R2, K4, the “center-tip” group includes residues R8, K9, K14, the “center-anchor” group includes residues R26 and K27, and the “anchor” group includes the residues anchoring the L-DNA to the inner gyre of the DNA K36, K37, R40 and R42. Both large opening events involved the loss of interactions between these residues with the outer DNA gyre.

**Figure 7:**
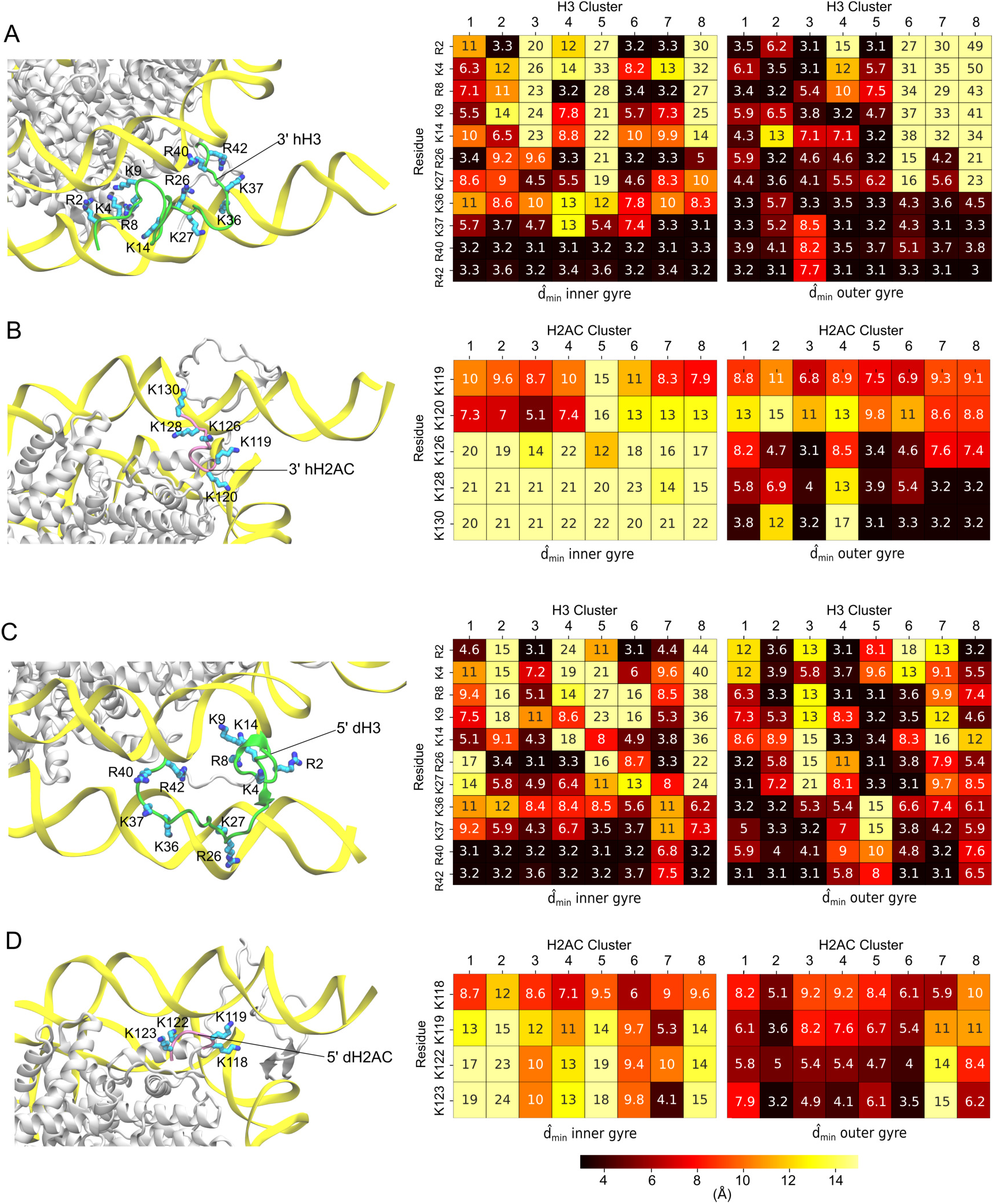
Interactions of epigenetic regulatory residues from the histone tails with DNA. Each panel shows a structural illustration of the histone tail (left) and 2 heatmaps with minimal distances of the analyzed residues to the inner (middle) and outer gyre of the DNA (right). **A-B**) the hH simulations. **C-D**) the dH simulations. The H3 (A,C) and H2AC (B,D) tails are shown as green and mauve ribbons with the analyzed aminoacids colored by atom name. The DNA is in yellow ribbons and the core histones are in white. The heatmaps show the analyzed residues on the vertical and the DNA clusters on the horizontal. The numbers are the median value of the minimal distance of each residue to the DNA in the corresponding DNA cluster.

In the open conformations of the Esrrb^hH^ nucleosome the H3 tail configurations(cluster *hH*3^6^) lacked interactions with the outer DNA gyre except for the hydrogen bonds formed by R42 from the anchor group. The interactions between the tip and center-tip groups and the outer DNA gyre were not formed in the simulation in which the nucleosome opened. The interactions of the center-anchor residues were lost before the nucleosome opened. Therefore, the loss of these interactions is a prerequisite for the opening (Supplementary Figure S6). Some of the residues from the tip and the center groups (R2, R8, R26) repositioned to interact with the inner gyre. Interestingly, most lysines involved in epigenetic regulation do not interact with DNA in these configurations (Figure 7A). The H2AC tail configurations in open nucleosomes (cluster *hH*2*AC*^4^) do not interact with DNA (Figure 7B), the interactions breaking before the opening. In contrast, in the closed conformations of Esrrb^hH^ K126, K128, and K130 formed stable interactions with the outer DNA gyre (Supplementary Figure S7).

In the open conformations of the Lin28b^dH^ nucleosome, the H3 tail configurations (cluster *dH*3^5^) interacts with the outer DNA gyre with the residues in the center groups (R8, K9, K14, R26, K27). The residues of the tip and anchor groups do not interact with the DNA while some anchor residues (K37, R40, R42) interact with the inner DNA gyre (Figure 7C, Supplementary Figure S6). In addition, the interactions between the dH2AC tail and the outer gyre broke before the opening (Figure 7D, Supplementary Figure S7).

In the hH nucleosome conformations open only in one direction (*γ*_1_ or *γ*_2_) the H3 tails main-tained some of its interactions with the outer DNA gyre. In the configurations from cluster *hH*3^3^ found in nucleosome conformations open in the XZ plane (*γ*_1_), the anchor group interacts with the inner DNA gyre (Figure 7A). On the other hand, the configurations from clusters *hH*3^2^, *hH*3^4^, and *hH*3^5^ found in nucleosome conformations open in the XY plane (*γ*_2_), the anchor group interacts with the outer DNA gyre, whereas the residues in the tip and center groups do not interact with DNA (Figure 7A).

In dH nucleosome conformations open in the XZ plane (γ_1_), the H3 configurations (clusters *dH*3^4^ and *dH*3^5^) have no interactions between the anchor group and the outer DNA gyre. On the other hand, in the configurations from clusters *dH*3^1^ and *dH*3^3^ found in nucleosomes open in XY plane (γ_2_), anchor but not tip and center residues interact with the outer gyre (Figure 7C).

In the closed nucleosome, the H3 and H2AC configurations have the most interactions with the outer DNA gyre. Therefore, the amplitude of nucleosome breathing depends on the number of interactions between aminoacids in the H3 and H2AC tails and the nearby L-DNA. When many of these are formed, the nucleosomes remained closed. For a large nucleosome opening, the loss of most but not all interactions between the tails and the L-DNA was required. The residues involved are located in clusters of positively charged residues and the interactions of one residue may be substituted by interactions of nearby residues from the same cluster. This suggests that these residues may be accessible for chemical modifications also in closed nucleosomes.

## Discussion

Chromatin fibers are not as compact as it has been thought (*57*). Hence, their folding and dynamics are more sensitive to the ratio between open and closed nucleosomes. It becomes evident that elucidating the mechanisms by which nucleosomes open and close is key to understand chromatin dynamics.

Here we showed how the interplay between the histone H3 and H2AC tails controls the breathing of genomic nucleosomes. From a total of 18 *μ*s of atomistic MD simulations, we observed 2 large opening events in 2 different genomic nucleosomes and only low amplitude breathing motions in an engineered nucleosome. Our findings suggest that there are lower barriers for large amplitude opening in genomic nucleosomes. This supports previous findings (*38*) that genomic nucleosomes are more flexible and mobile than artificial nucleosomes bearing strong positioning DNA sequences such as the Widom 601 sequence (*39*).

Previously, we reported that the larger structural flexibility of the Lin28b nucleosome with *Drosophila* histones was in agreement with its lower thermal stability in experiments (*38*). Lin28b opened extensively in the simulations with Drosophila histones and adopted more open conformations in all simulations, suggesting that it is the less compact nucleosome.

Here we also report a high structural flexibility leading to a large opening event in the Esrrb nucleosome with human histones in contrast to the Esrrb nucleosome with Drosophila histones which was more rigid. The Esrrb nucleosome had a lower thermal stability than the Widom nucleosome but higher than Lin28. Moreover, its stability varied in the experiment (*38*), suggesting that it may adopt both more or less stable conformations.

However, caution is needed when comparing the simulations with these experiments. Although both nucleosome disassembly in the experiments and nucleosome breathing in the simulations are measures of nucleosome structural flexibility, they differ in amplitude and time scale of the motions involved. We propose that the variability in our simulations is not due to the minor differences between the human and Drosophila histone sequences but rather to the differential sampling of histone tail dynamics (see discussion below).

Complete nucleosome unwrapping is thought to happen in seconds, with intermediate states forming after hundreds of milliseconds (*58,59*). While atomistic MD simulations are very powerful in in studying nucleosome dynamics, the timescales accessible are in the *μ*s range (*35,36,38,60*). Therefore, the simulations are not converged and achieve only a limited sampling of the conformational space. This limitation may be circumvented by enhanced sampling techniques (*60, 61*). However, these involve biasing the motions which may provide inaccurate estimations of the energies and pathways of nucleosome motions (*60*). We argue that the grouping of multiple independent simulations started from the same structure (replicas) in ensembles is a powerful approach to achieve extensive sampling. In each replica, the nucleosome samples a subregion of the phase space and two replicas are never identical because of the randomizations performed at the beginning of each simulation. Therefore, merging the replicas expands the phase space explored. This approach enabled us to monitor the 2 rare large opening events in complete nucleosomes and study their mechanisms. The *γ*_1_-γ_2_ histograms demonstrated that the openings were extensions of the low amplitude nucleosome breathing. In contrast to previous reports of larger amplitude opening in atomistic simulations of incomplete nucleosomes, we could elucidate the role of histone tails in these motions (*62*).

Histone tails are important for chromatin dynamics. H3 and H4 tails impact the packing of the genome by modulating the interaction between nucleosomes (*6, 7*). Moreover, the tails interfere with the binding of other proteins to the nucleosome. For example, the H2AC tail interacts with the linker histone impacting on its role to increase chromatin compaction upon binding to nucleosomes (*8*). However, the mechanisms by which the tails impact on nucleosome structural flexibility remained obscure.

The role of the tails has been proposed from a number of coarse grained simulations (*15,18, 21,23*). In these simulations, the histone tails are simplified and the detailed chemistry of the interactions involved is often neglected, leading to an incomplete representation of the protein-DNA interactions. Precisely this gap is closed by atomistic simulations. However, simulating large amplitude nucleosome motions and accurately describing the dynamics of histone tails is challenging. The simulations of intrinsically disordered proteins or protein regions like histone tails is particularly difficult, because the force-fields used for structured proteins often do not provide a correct sampling of their conformational space (*63*). However, the force fields we used represent the state-of-the-art, and reproduce the dynamics of intrinsically disordered proteins with reasonable accuracy (*63*)). Force fields from this family have been used to explain the destabilization of the nucleosome by mutations in H2A in experiments, confirming their accuracy (*64*).

The atomistic studies to date have been performed using either incomplete nucleosome or nucleosomes with artificial sequences such as the Widom or the human *α*-satellite DNA. Large scale breathing observed in tail-less nucleosomes provided insights into the interactions of the histone core with DNA. However, removing the histone tails in these simulations lead to unnatural breathing (*24,26–29*).

Here, we report a regulation mechanism for breathing of complete, genomic nucleosomes that involves an interplay between the histone H3 and H2AC tails and specific interactions between residues in these tails and the DNA. The general behavior of the H3 and H2AC tails was similar in the simulations with *Drosophila* and human histones. The tails collapsed on the DNA and large changes in the tail conformation, position, and DNA binding pattern were rare in 1 *μ*s. However, the tail configurations were different in the simulations of the same ensemble, allowing a more extensive sampling. Our finding explain previous reports that the deletion of either of the 2 tails lead to large amplitude nucleosome opening in experiments (*8, 65, 66*) and in high temperature simulations (*37*). We propose that the tails keep nucleosome mostly closed and a cooperative motion of the H3 and H2AC tails is needed for a large amplitude opening.

Post-translational, chemical modifications of key residues in histone tails mark chromatin regions as active or inactive, establishing the epigenetic regulation of gene expression at a given time and cellular context. Therefore, it is of utmost importance to understand the role of these residues in nucleosome and chromatin dynamics. We show that epigenetically active residues in the H3 tail are located in clusters of positively charged residues that interact with DNA. We observed that for a large opening of nucleosomes, the majority of these interactions, especially those with the L-DNA have to be absent. Our findings suggest that the nucleosome is kept closed when a majority of these interactions are formed. However, the absence of interactions of each individual residue may be compensated by interactions of residues from the same cluster, suggesting that these residues may be available for chemical modification even in closed nucleosomes. For example, K9 and K27, both methylated in inactive chromatin did not establish specific interaction with DNA independent in any nucleosome conformation. However, arginines flanking them displayed a nucleosome conformation dependent pattern of interactions with DNA. K36 from the region anchoring L-DNA to the core DNA is acetylated in active chromatin and formed interactions with the DNA only in closed nucleosomes, suggesting that this residue is more accessible in open nucleosomes. However, K36 is flanked by 2 arginines and another lysine forming redundant interactions with the DNA. Therefore, our findings suggest the tail residues are accessible for chemical modification independent of nucleosome conformation. The impact of such modifications on intra-nucleosome motions may manifest in more subtle ways than just by neutralizing the positive charge of the tail residues. Studying how single modifications or histone variants affect nucleosome dynamics with atomistic MD simulations is a powerful alternative to experiments (*36,61,67,68*). We predict that with the continuous increase of computational resources available, long simulations of complete genomic nucleosomes, modified or unmodified will contribute to elucidating these mechanisms

## Acknowledgements

The authors thank Caitlin M MacCarthy for the support and discussion. J.H. is part of the International Max Planck Research School-Molecular Biomedicine, Münster, Germany. This work was supported by funds of the Max Planck Society and The Royal Netherlands Academy of Arts and Sciences. Computer resources were provided by the Gauss Centre for Supercomputing e.V. (www.gauss-centre.eu) (project ID 12622, STRUCNUCREC running on the GCS Supercomputer SuperMUC at the Leibniz Supercomputing Centre (www.lrz.de), to J.H. and V.C.).

## Supplementary Materials

Supplementary Files Huertas et al.pdf: PDF file containing Supplementary Documents S1 and S2, and Supplementary Figures S1 to S7

Supplementary Document S3.xlsx: Excel file containing all the distances and contacts of the H3 and H2AC tails, over time, in all simulations.

