## Supplementary Data for "Histone tails cooperate to control the breathing of genomic nucleosomes"

#### Document S1. Sequences of Widom, Esrrb and Lin28B used in the simulations.

##### Widom

```
  1 ACGCGGCCGC CCTGGAGAAT CCCGGTGCCG AGGCCGCTCA ATTGGTCGTA
 51 GCAAGCTCTA GCACCGCTTA AACGCACGTA CGCGCTGTCC CCCGCGTTTT
101 AACCGCCAAG GGGATTACTC CCTAGTCTCC AGGCACGTGT CAGATATATA
151 CATCCTGTGC ATGTATTG
```

##### Esrrb

```
  1 ATCAGCAGGG AGAAGGAGCG CCTCCCCATG TGGGACCTGG AGAAACAGAG
 51 GGTGGAGGGA GCATAGAGAG TCTGTTCTAA GCTGCAAAGC AAAGGCCTGG
101 CGACCTAGGA GACCATGGAG TTCCAGAAAG TGATAGTTAT GCAGAGCGAA
151 TGGAGGGAAT CAGCACGC
```

##### Lin28b – Drosophila histones simulations

```
  1 AGTTAAGTGG TATTAACATA TCCTCAGTGG TGAGTATTAA CATGGAACTT
 51 ACTCCAACAA TACAGATGCT GAATAAATGT AGTCTAAGTG AAGGAAGAAG
101 GAAAGGTGGG AGCTGCCATC ACTCAGAATT GTCCAGCAGG GATTGTGCAA
151 GCTTGTGAAT AAAGACAC
```

##### Lin28b – Human histones simulations

```
  1 AAGTTAAGTG GTATTAACAT ATCCTCAGTG GTGAGTATTA ACATGGAACT
 51 TACTCCAACA ATACAGATGC TGAATAAATG TAGTCTAAGT GAAGGAAGAA
101 GGAAAGGTGG GAGCTGCCAT CACTCAGAAT TGTCCAGCAG GGATTGTGCA
151 AGCTTGTGAA TAAAGACA
```

### Document S2. Histone Alignments

## H3

```
                          N-terminal tail
Human        MARTKQTARKSTGGKAPRKQLATKAARKSAPATGGVKKPHRYRPG
Drosophila   MARTKQTARKSTGGKAPRKQLATKAARKSAPATGGVKKPHRYRPG

Human        TVALREIRRYQKSTELLIRKLPFQRLVREIAQDFKTDLRFQSSAV
Drosophila   TVALREIRRYQKSTELLIRKLPFQRLVREIAQDFKTDLRFQSSAV

Human        MALQEACEAYLVGLFEDTNLCAIHAKRVTIMPKDIQLARRIRGER
Drosophila   MALQEASEAYLVGLFEDTNLCAIHAKRVTIMPKDIQLARRIRGER

Human        A
Drosophila   A
```

## H4

```
                N-terminal tail
Human        MSGRGKGGKGLGKGGAKRHRKVLRDNIQGITKPAIRRLARRGGVK
Drosophila   MTGRGKGGKGLGKGGAKRHRKVLRDNIQGITKPAIRRLARRGGVK

Human        RISGLIYEETRGVLKVFLENVIRDAVTYTEHAKRKTVTAMDVVYA
Drosophila   RISGLIYEETRGVLKVFLENVIRDAVTYTEHAKRKTVTAMDVVYA

Human        LKRQGRTLYGFGG
Drosophila   LKRQGRTLYGFGG
```

## H2A

*N-terminal tail*

```
Human   MSGRGKQGGKARAKAKTRSSRAGLQFPVGRVHRLLRKGNYSERVG
Droso   MSGRGK.GGKVKGKAKSRSNRAGLQFPVGRIHRLLRKGNYAERVG

Human   AGAPVYLAAVLEYLTAEILELAGNAARDNKKTRIIPRHLQLAIRN
Droso   AGAPVYLAAVMEYLAAEVLELAGNAARDNKKTRIIPRHLQLAIRN
```

*C-terminal tail*

```
Human   DEELNKLLGRVTIAQGGVLPNIQAVLLPKKTESHHKAKGK
Droso   DEELNKLLSGVTIAQGGVLPNIQAVLLPKKTEKKA.....
```

## H2B

*N-terminal tail*

```
Human   MPEPAKSAPAPKKGSKKAVTKAQKKDGKKRKRSRKESYSVYVYKV
Droso   MP.PKTSGKAAKKAG.KAQKNITKTD.KKKKRKRKESYAIYIYKV

Human   LKQVHPDTGISSKAMGIMNSFVNDIFERIAGEASRLAHYNKRSTI
Droso   LKQVHPDTGISSKAMSIMNSFVNDIFERIAAEASRLAHYNKRSTI

Human   TSREIQTAVRLLLPGELAKHAVSEGTKAVTKYTSSK
Droso   TSREIQTAVRLLLPGELAKHAVSEGTKAVTKYTSSK
```

**Document S3. Position of the H3 and H2AC tails.** xlsx file with the distances and contacts of the H3 and H2AC tails, over time, in all simulations.

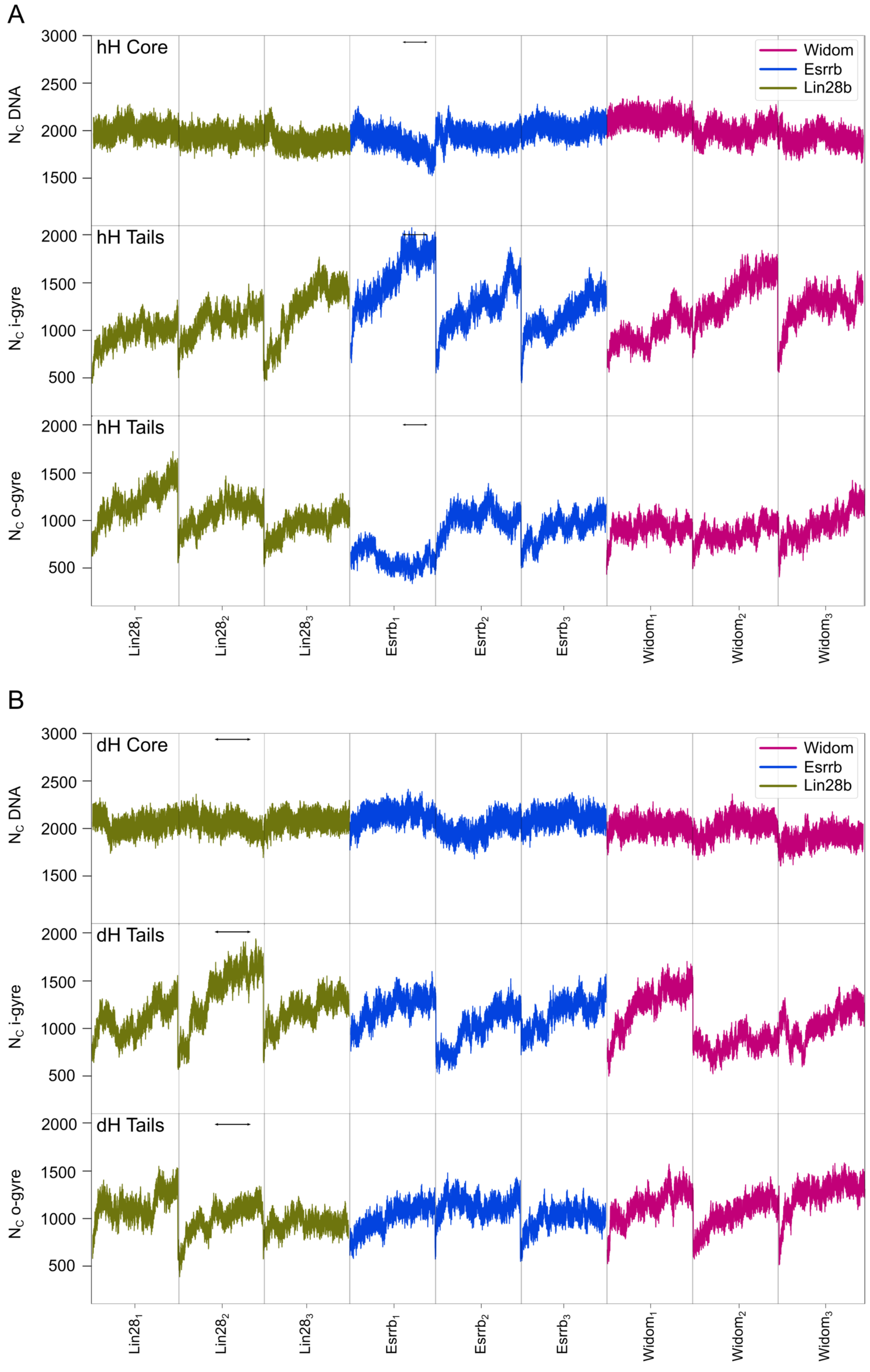


Supplementary Figure S1: **Number of histone-DNA contacts.** The 2 panels depict the time evolution of the number of contacts of the histone core (top) and histone tails (middle and bottom) to the DNA. For the histone tails, contacts were split between contacts to the inner gyre (middle) and contacts to the outer gyre (bottom). **A)** Contact evolution in the hH simulations. **B)** Contact evolution in the dH simulations. A contact was defined as a non-hydrogem atom closer than 4.5 Å to another non-hydrogen atom.

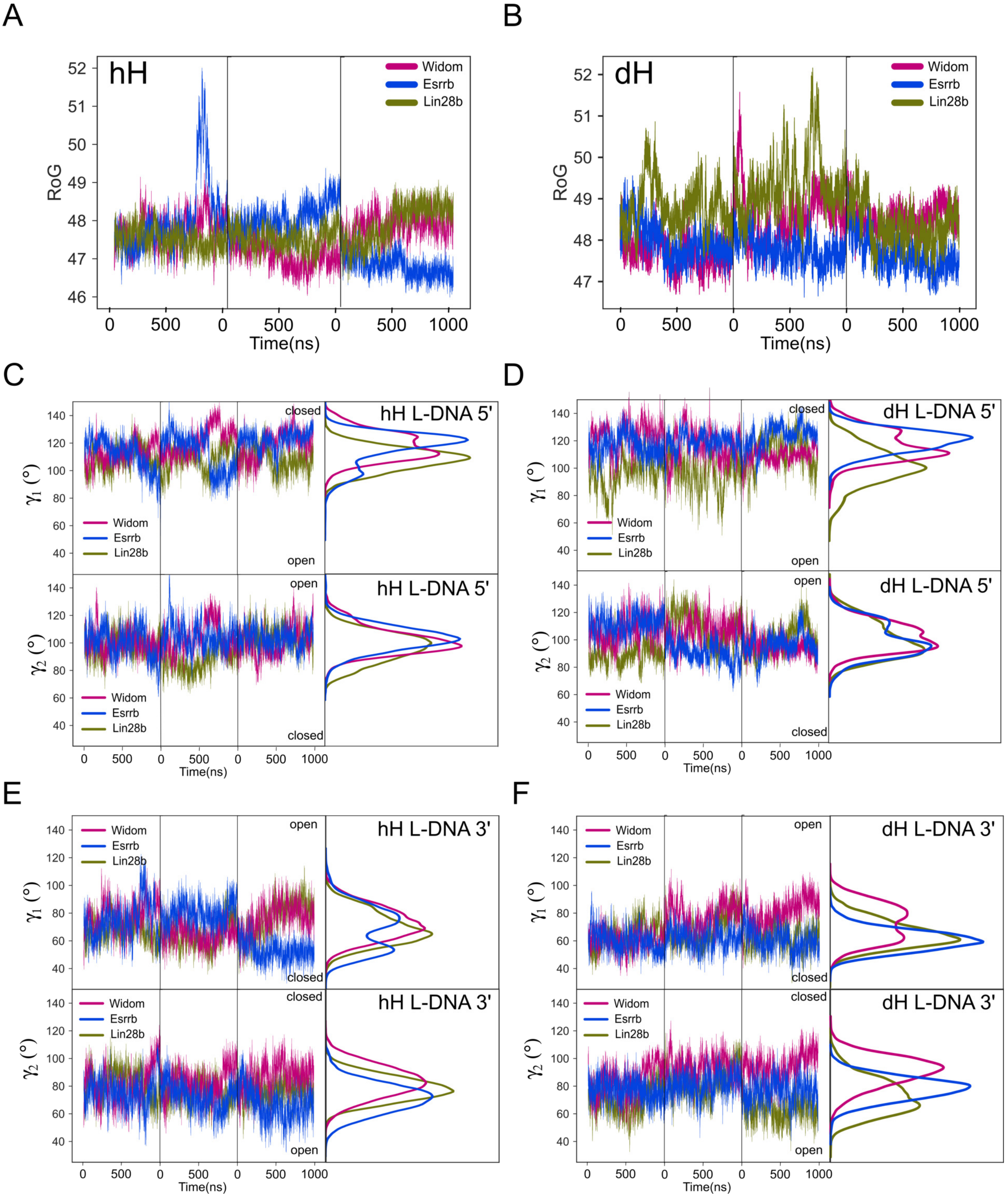


Supplementary Figure S2: **Nucleosome structural flexibility A-B)**: The evolution of the RoG of the DNA over time, during the hH (A) and dH (B) simulations. **C-F)**: Time series and histograms of the $\gamma$ angles described in Figure 2, for the 5' arm of the hH nucleosomes (C), 5' arm of the dH nucleosomes (D), 3' arm of the hH nucleosomes (E) and 3' arm of the dH nucleosomes (F). The individual simulations are separated by vertical black lines.

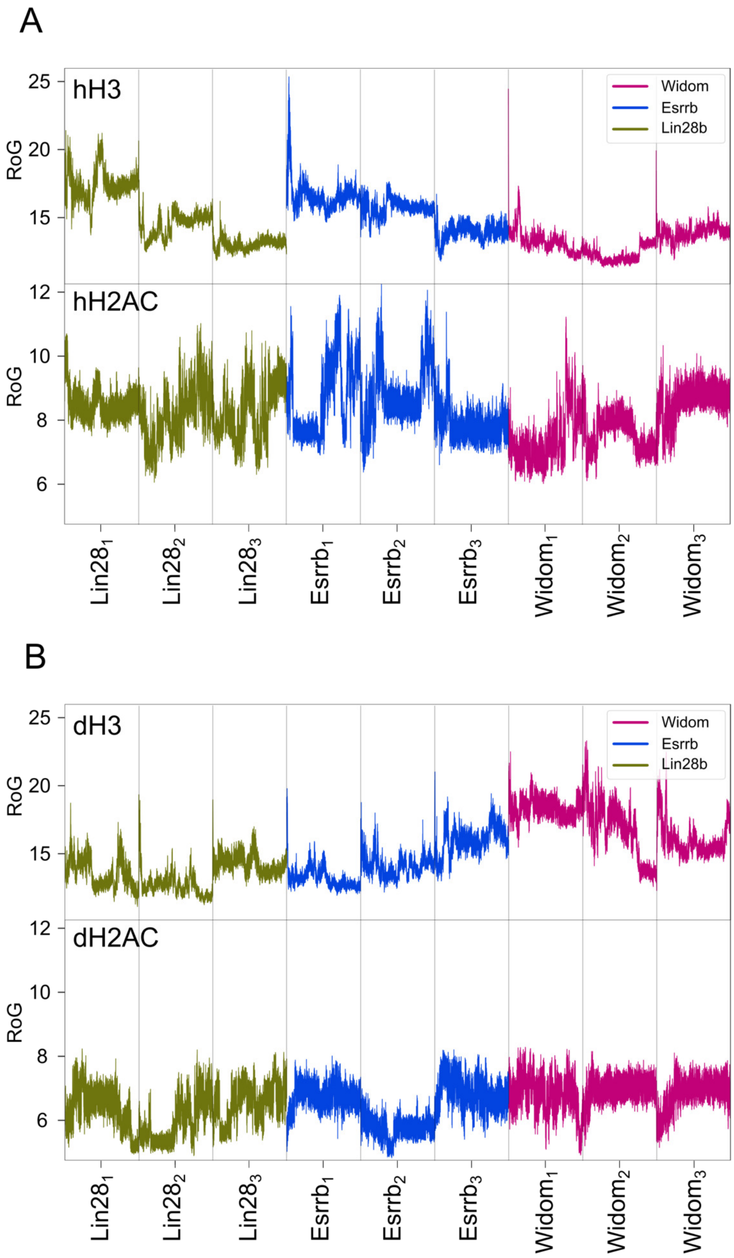


Supplementary Figure S3: **Radius of gyration of the histone tails** The evolution of the RoG over time for histones H3 and H2AC tails. **A)** hH simulations. **B)** dH simulations.

### A. hH DNA 3'

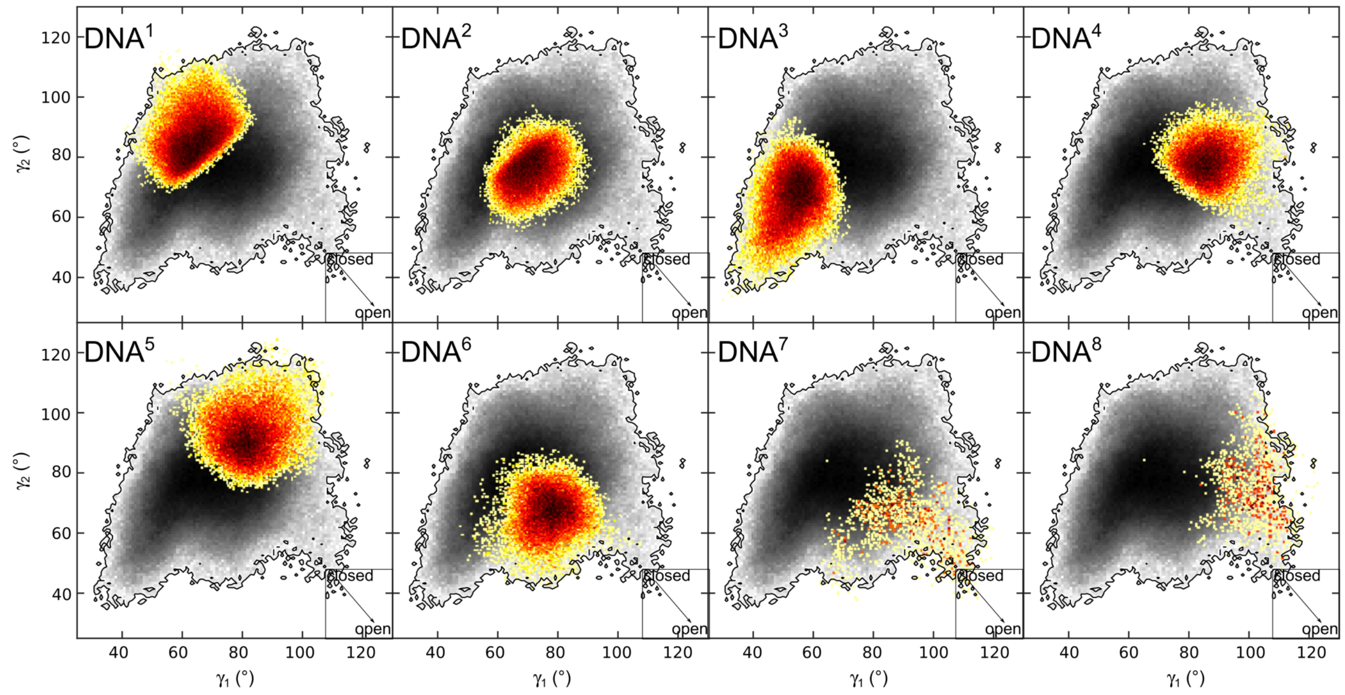


### B. dH DNA 5'

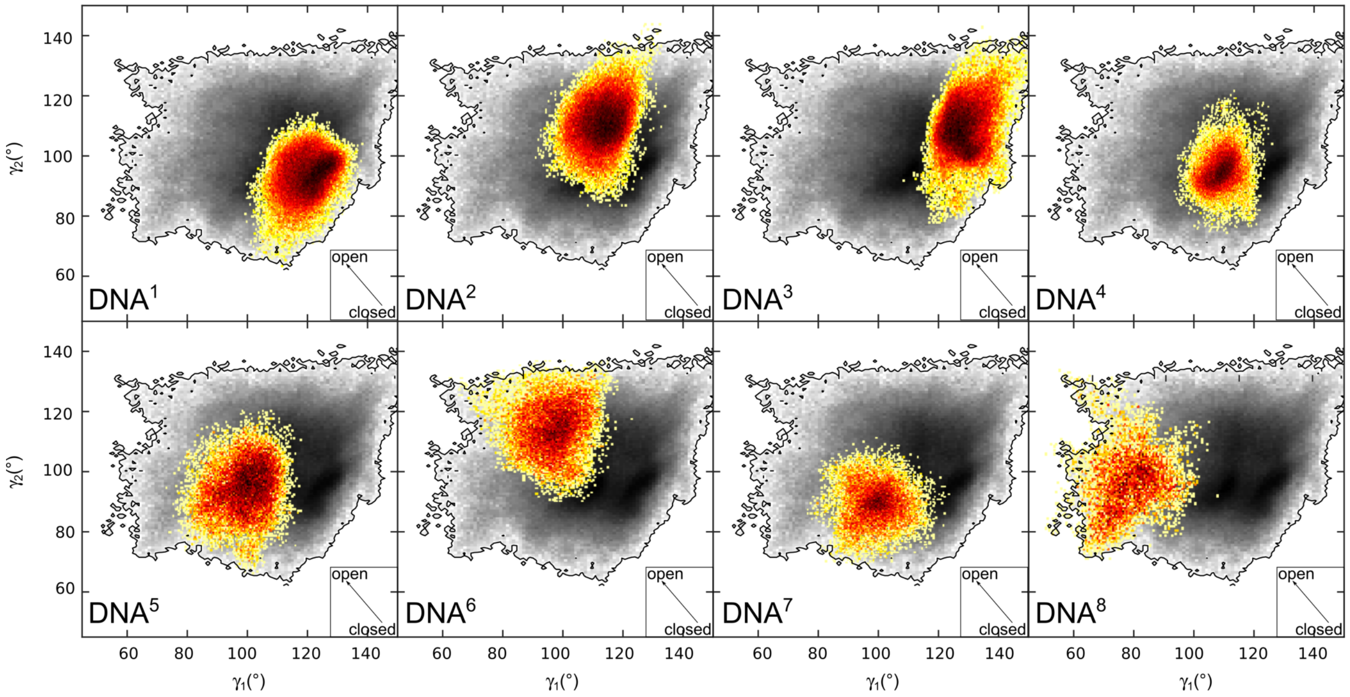


Supplementary Figure S4: **Sampling of the DNA clusters** Two-dimensional histograms depicting the conformational sampling of the L-DNA arms in the space defined by the $\gamma_1$ and $\gamma_2$ angles for each individual DNA cluster. **A)** 3' L-DNA clusters from the hH simulations. **B)** 5' L-DNA clusters from the dH simulations

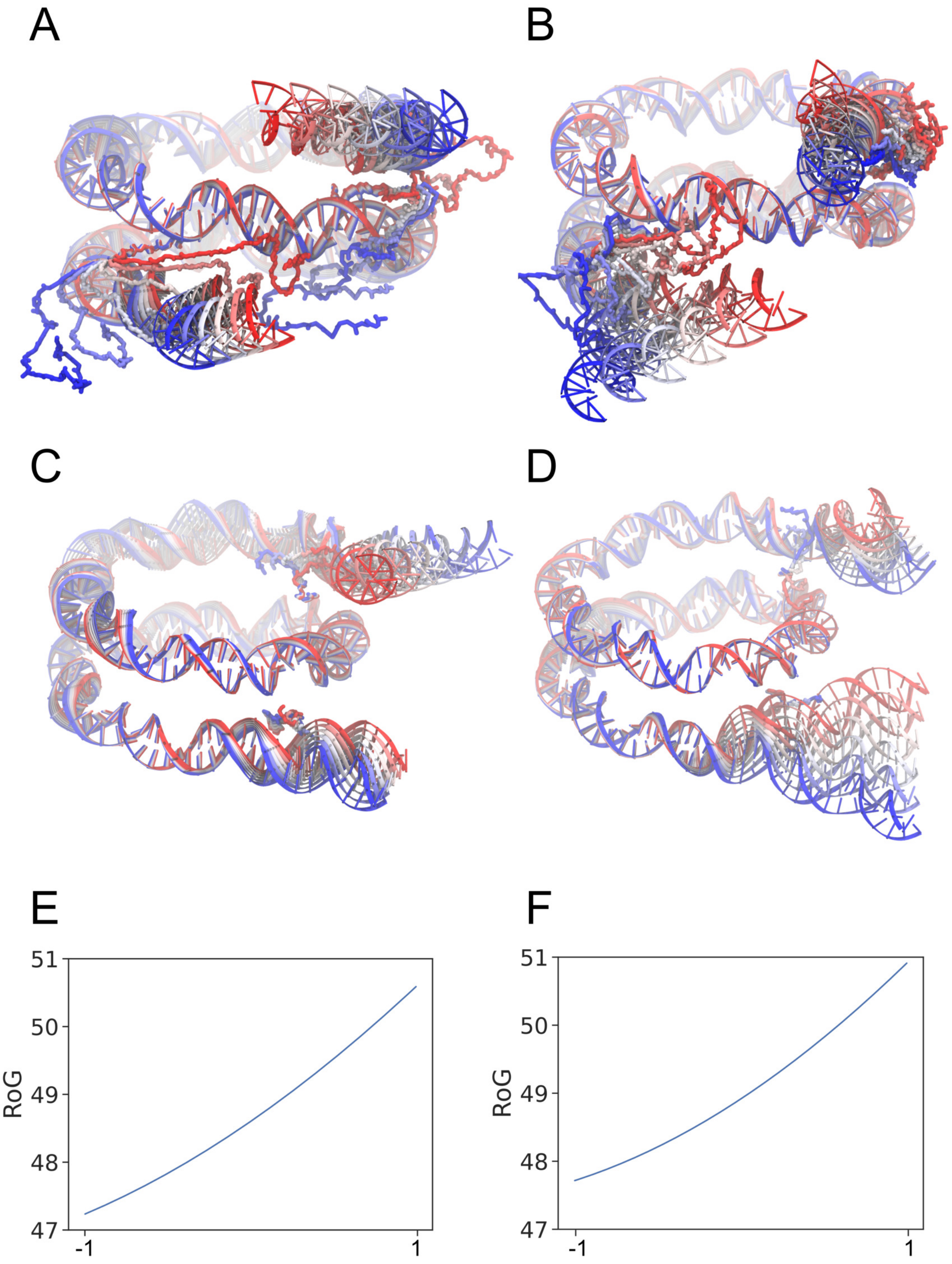


Supplementary Figure S5: **Correlated motions of the histone tails and linker DNA. A-D)** Superposition of DNA and H3 (A-B) or DNA and H2AC (C-D) snapshots from the pseudotrajectory of the lowest frequency principal component (PC1) of the simulation ensembles of Esrrb$^{\mathrm{hH}}$ (A,C) and Lin28b$^{\mathrm{dH}}$ (B,D). **E-F)** The RoG along PC1 pseudotrajectory of Esrrb$^{\mathrm{hH}}$ (E) and Lin28b$^{\mathrm{dH}}$ (F). The closed to open transition is indicated by the Red-White-Blue color scale. The amplitude was normalized such as -1 corresponds to the most closed and +1 to the most open conformation.

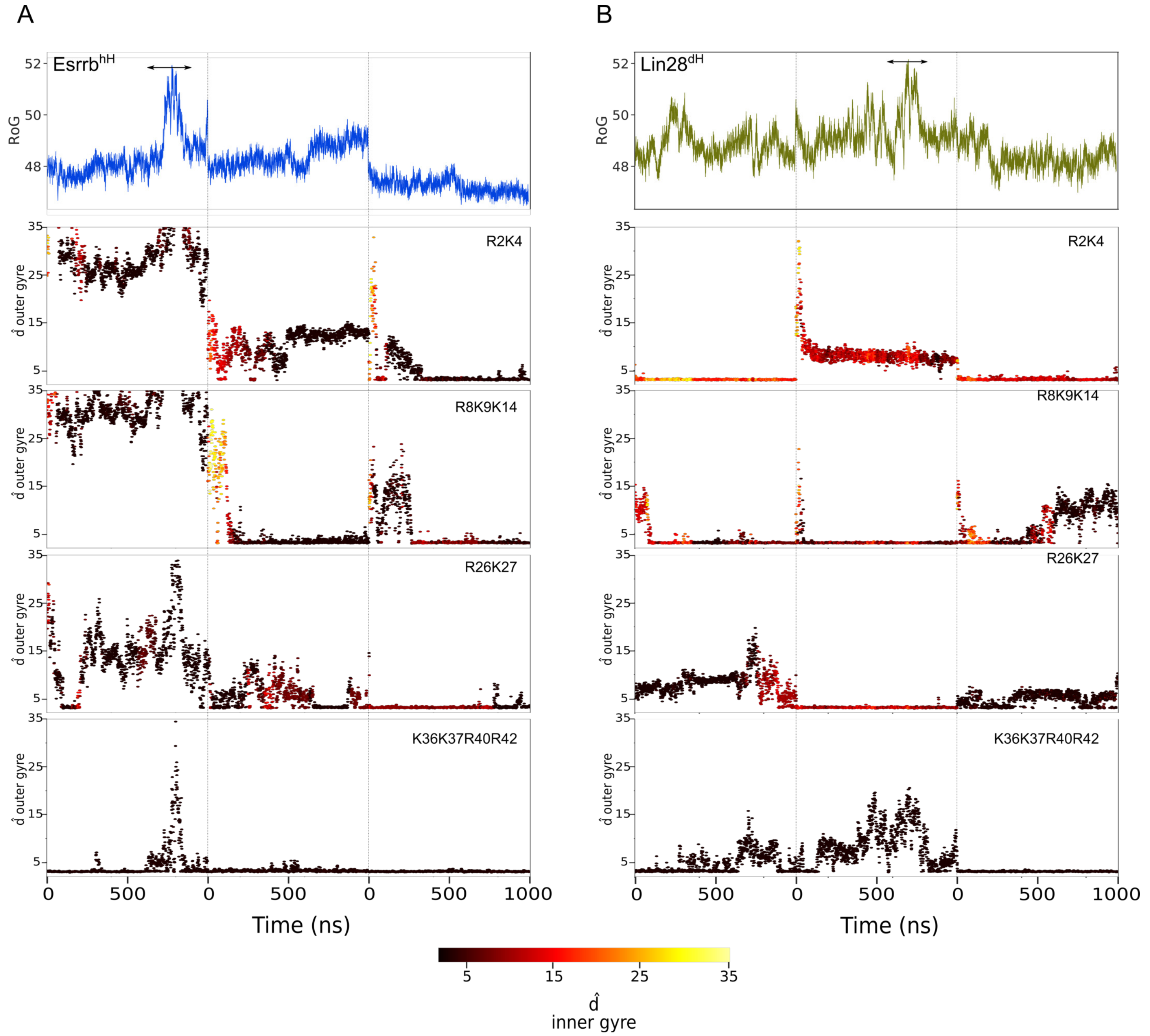


Supplementary Figure S6: **Interactions of H3 residues with DNA.** The evolution of the H3 residues position relative to the inner and outer gyre of the DNA. **A)** The Esrrb$^{hH}$ nucleosome. **B)** The Lin28b$^{dH}$ nucleosome. The plot at the top row shows the nucleosome RoG to monitor opening and closing events. The other plots show the minimal distance of the residues to the outer gyre, colored by the minimal distance to the inner gyre of the DNA.

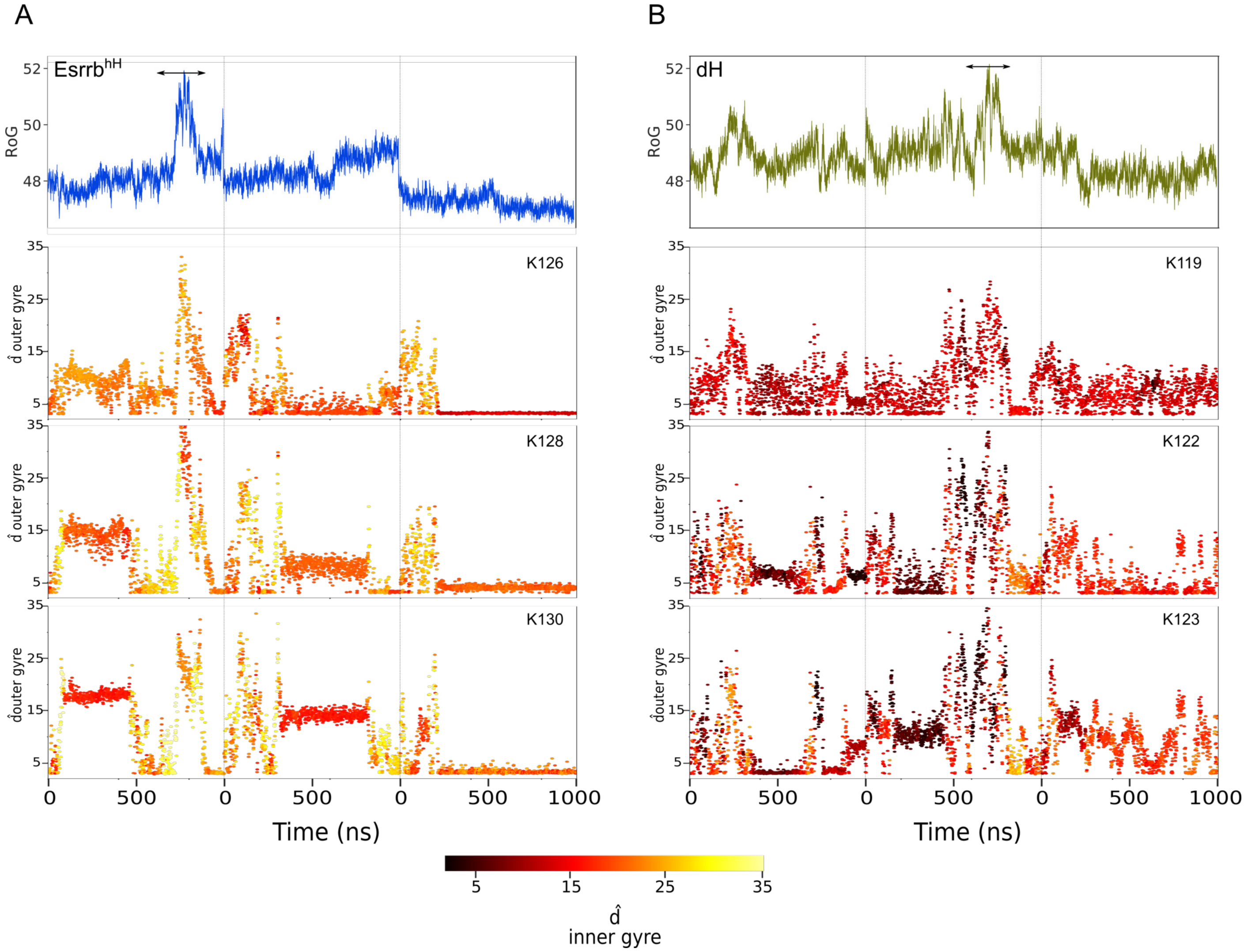


Supplementary Figure S7: **Interactions of H2AC residues with DNA in the nucleosomes with large opening.** The evolution of the H2AC residues position relative to the inner and outer gyre. **A)** The Esrrb$^{hH}$ nucleosome. **B)** The Lin28b$^{dH}$ nucleosome. The plot at the top row shows the nucleosome radius of gyration to monitor opening and closing events. The other plots show the minimal distance of the residues to the outer gyre, colored by the minimal distance to the inner gyre of the DNA.
